# Type I interferons sensitise HIV-1-reactivating T-cells for NK cell-mediated elimination despite HDACi-imposed dysregulation of innate immunity

**DOI:** 10.1101/2020.05.04.075119

**Authors:** Julia Prigann, Dylan Postmus, Anna Julia Pietrobon, Emanuel Wyler, Jenny Jansen, Lars Möller, Jelizaveta Fadejeva, Thijs H. Steijaert, Cornelius Fischer, Uwe Koppe, Barbara Gunsenheimer-Bartmeyer, Karolin Meixenberger, Sarah N. Vitcetz, Madlen Sohn, Lucie Loyal, Andreas Thiel, Sascha Sauer, Kathrin Sutter, Ulf Dittmer, Michael Laue, Norbert Bannert, Markus Landthaler, Christine Goffinet

## Abstract

Shock-and-kill is one of the most advanced, yet unrealized, concepts towards establishment of HIV-1 cure. Treatment with latency-reversing agents (LRAs), including histone deacetylase inhibitors (HDACis) exerting chromatin remodelling and gene expression reprogramming, combined with anti-retroviral therapy reactivates HIV-1 transcription *in vitro*, *ex vivo* and *in vivo*. However, HDACi treatment fails to significantly reduce the size of the viral reservoir in people living with HIV-1 (PLHIV). Here, by combining scRNA-seq and functional approaches, we characterised the HDACi treatment-imposed remodulation of CD4+ T-cells’ state and its consequences for HIV-1 latency reversal and the apparent resistance of HIV-1-reactivating cells to immune-mediated elimination. Exposure of CD4^+^ T-cells from three aviremic PLHIV with clinically applicable concentrations of Panobinostat markedly reduced the expression of genes mediating T-cell activation and IFN-driven antiviral immunity in a largely CD4^+^ T-cell subset-nonspecific manner, with exception of an PLHIV-specific exhausted CD4^+^ T-cell subpopulation. Altered transcriptomic profiles were accompanied by large refractoriness to peptide and IL-2/PHA stimulation, and to exogenous type I interferon, that would otherwise induce T-cell activation and expression of a plethora of antiviral genes, respectively. Type I interferon, when added to Panobinostat during HIV-1 reactivation, was unable to counteract HDACi-mediated inhibition of IFN signalling and failed to interfere with HIV-1 reactivation *per se*. However, it imposed a pre-budding block and boosted surface levels of HIV-1 Env on reactivating cells. Co-treatment with type I IFNs, most prominently IFN-β and -α14, sensitised HIV-1-reactivating cells for killing by NK cells through antibody-dependent cytotoxicity. Together, our study provides proof-of-concept of the benefit of combining a potent LRA with immunostimulatory molecules, such as type I IFNs, to reduce the resistance of HIV-1-reactivating T-cells to immune-mediated elimination to improve current shock-and-kill strategies.

## INTRODUCTION

Antiretroviral therapy (ART) effectively suppresses viremia and has improved the quality and duration of life of people living with HIV/AIDS (PLHIV), but requires livelong adherence and is not curative (1–4). HIV-1 persists as a viral reservoir in latently infected, long-lived CD4+ T-cells with an estimated 73 years of optimal ART theoretically required for reservoir eradication (5,6).

To accomplish ART-free remission, studies have focused on the reduction of the size of the latent reservoir to achieve a functional HIV-1 cure (7). The shock-and-kill cure approach is based on the administration of latency-reversing agents (LRAs) combined with ART with the intention to reverse proviral quiescence in cellular reservoirs, an event that should be followed by specific elimination of HIV-1-reactivating cells using immunological or pharmacological mechanisms (8,9). Various classes of molecules, such as histone deacetylase inhibitors (HDACis) (10–12), protein kinase C (PKC) agonists (13–15), bromodomain and extraterminal domain (BET) protein inhibitors (16), second mitochondria-derived activator of caspases (SMAC) mimetics (17), and Toll-like receptor (TLR) agonists (18,19) have been explored for their ability to reverse HIV-1 latency *ex vivo* and *in vivo.* In clinical trials individual LRAs demonstrate clear ability to reactivate HIV-1 transcription, demonstrated by increased plasma viremia during continued ART. However, the size of the viral reservoir in most studies decreased only to a minor extent, if at all (8,20) for poorly understood reasons. One potential exception is the recent study in which treatment with Venetoclax, a pro-apoptotic inhibitor of BCL-2, delayed viral rebound in a humanised mouse model of HIV-1 infection, and depleted integrated HIV-1 DNA in CD4^+^ T-cells from PLHIV (21), suggesting that sensitising HIV-1-positive cells to apoptosis might enable reduction of the reservoir size. Overall, existing data demonstrate an urgent need for a conceptual improvement of the shock-and-kill approach.

Besides CD8^+^ T-cell mediated mechanisms (22), NK cell-mediated antibody-dependent cellular cytotoxicity (ADCC) has been proposed to contribute to natural, ART-free control of HIV infection *in vivo*. Specifically, potent NK cells’ anti-HIV activity was identified as a discriminatory parameter in HIV-1 elite controllers as opposed to viremic progressors (23,24) and as a specific characteristic of the few participants in the RV144 HIV-1 vaccine trial who were protected from infection after immunisation (25,26). Type I IFNs increase the potency of NK cells to eliminate HIV-1-infected cells *ex vivo* (27,28). Administration of pegylated IFN-α2a or -α2b in combination with ART has resulted in sustained viral control in PLHIV and declined levels of integrated HIV-1 DNA in the context of structured ART interruptions, observations that associated with enhanced cytotoxic NK cell activity (29–35). Overall, there is a growing body of evidence pointing towards a potential benefit of incorporating type I IFNs into LRA-involving HIV-1 cure strategies, although the individual combinations of specific LRAs with type I IFN have not yet been functionally tested in regards to both HIV-1 transcriptional reactivation and cell elimination.

Here, we characterise the transcriptional and functional profiles of CD4^+^ T-cells from PLHIV upon *ex vivo* exposure to two HDACis, Panobinostat and Vorinostat, to explore potential cell-intrinsic properties contributing to the resistance of HIV-1-reactivating cells to immune-mediated elimination *in vivo*. Using single-cell RNA sequencing we identified *ex vivo* HDACi treatment to highly impact T-cell receptor and IFN-related innate immune signalling pathways, resulting in a strong impairment of respective functions. Despite the extensive shut-down of IFN signalling by Panobinostat treatment, type I IFN treatment, while not interfering with HIV-1 reactivation *per se*, induced a block prior to viral budding. This phenotype was accompanied by accumulation of viral Env on the surface of HIV-1-reactivating cells which translated into a higher susceptibility to NK cell-mediated ADCC as compared to cells treated with HDACi alone. In conclusion, this study provides proof-of-concept for combination of potent LRAs with type I IFNs being a promising strategy to overcome the resistance of HIV-1-reactivating T-cells to NK cell-mediated elimination. We propose that the results of our study have important implications for improving current shock-and-kill strategies and will inform future HIV-1 cure therapies.

## RESULTS

### Panobinostat treatment modulates expression of CD4^+^ T-cell subset-specific markers

In order to understand LRA-induced cellular phenotypes and CD4^+^ T-cell subpopulation-specific susceptibilities to HDAC inhibition, we exposed purified CD4^+^ T-cells isolated from three aviremic PLHIV (**Supplementary Table 1**) to Vorinostat, Panobinostat, IL-2/PHA or left them mock-treated for 48 hours, followed by single cell RNA-sequencing. We selected Vorinostat and Panobinostat as prototypic HDACis due to their demonstrated ability to reactivate HIV-1 RNA expression *in vivo* (36,37). The applied concentrations of Vorinostat (500 nM) and Panobinostat (50 nM) approached those detected in patientś plasma after single oral administration (11,36–38). We included IL-2/PHA treatment as a reference that we expected to result in maximal activation of T-cells.

We used previously defined marker genes to identify the following T-cell subsets (39–41): Naïve (T_N_), central memory (T_CM_), transitory memory (T_TM_), effector memory (T_EM_), effector memory re-expressing CD45RA (T_EMRA_) and regulatory (T_REG_) T-cells in the dataset of mock-, HDACi- and IL-2/PHA-treated CD4^+^ T-cells of PLHIV (**Fig. 1A**). In line with previous studies (42), the T-cell subset distribution was not significantly altered in mock-treated samples from aviremic PLHIV undergoing ART as compared to a culture from an HIV-1-negative donor (**Fig. 1B**). However, we confirmed the previously reported presence of a small subpopulation specifically identified in cells from PLHIV (42,43), characterised by a combined and significantly higher expression of the exhaustion markers *PDCD1*, *LAG3, HAVCR2, GZMB* that was clearly separated from the otherwise similar T_EMRA_ cells (**Sup. Fig. 1**). Accordingly, we referred to this subset as exhausted T-cells (T_EX_).

**Figure 1.**
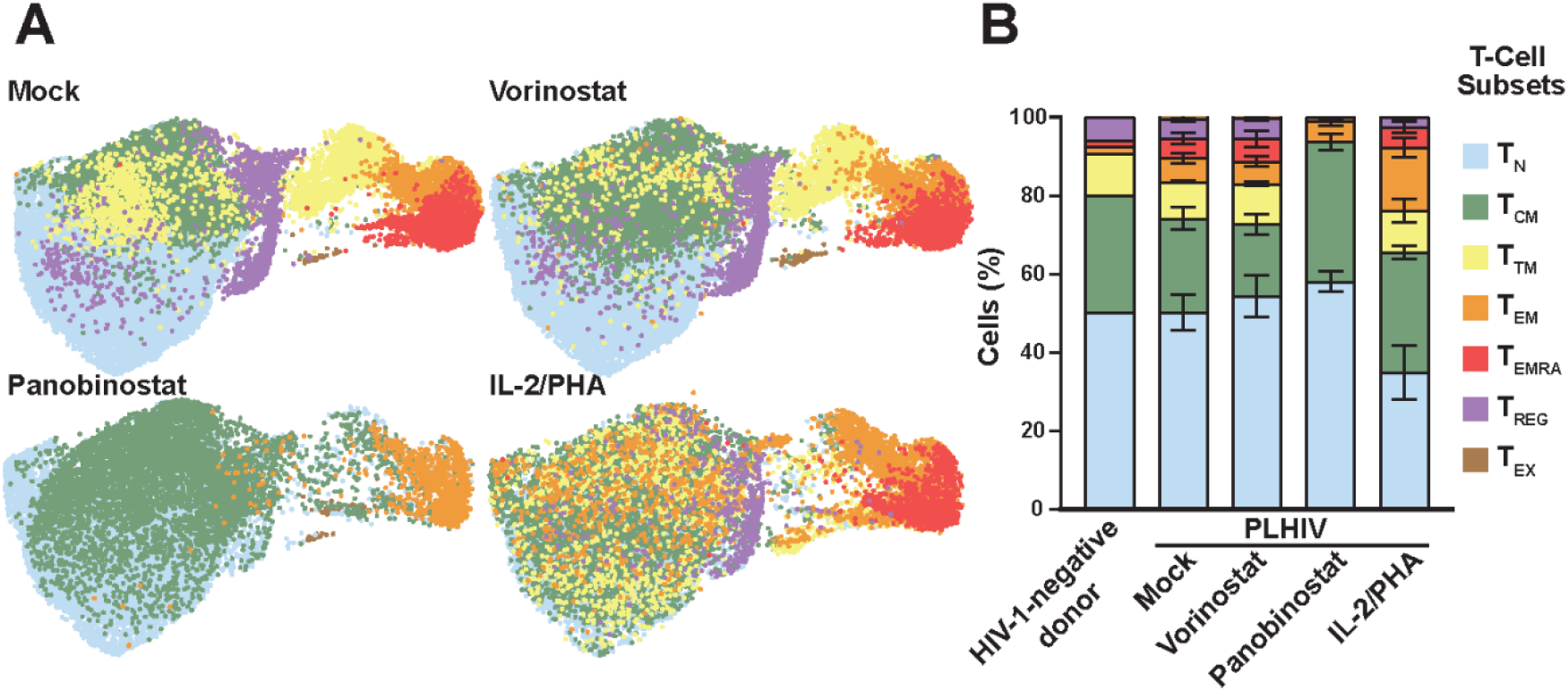
Panobinostat treatment modulates expression of CD4^+^ T-cell subset-specific markers. **(A)** Combined UMAP plot of CD4^+^ T-cell cultures from three PLHIV, separated by treatment and coloured by T-cell subset. **(B)** Percentages of indicated CD4^+^ T-cell subsets.

Overall, Vorinostat-treated CD4^+^ T-cell cultures displayed very mild changes in subset distribution as opposed to Panobinostat-treated culture that experience a complete loss of cells expressing markers of T_TM_ (9.34% to 0%, p = 0.0008), T_EMRA_ (5.03% to 0%, p = 0.0676) and T_REG_ (4.98% to 0%, p = 0.0161) subsets and a parallel increase in the proportion of T_CM_ (23.9% to 35.57%, p = 0.0155). Relative levels of T_N_ (50.25% to 54.34%, p = 0.0821), T_EX_ (0.43% to 1.11%, p = 0.778) and T_EM_ cells (6.06% to 5.18%, p = 0.2215) remained unchanged under these experimental conditions (**Fig. 1B**). The observed changes in the T-cell subset distribution induced by the individual treatments were detected in cell cultures from all three donors to a similar extent (**Sup. Fig. 2**). Although we confirmed a low, but statistical significant increase in both early and late apoptotic T-cells following Panobinostat treatment using a flow cytometry-base viability assay, the complete loss of T_TM_, T_EMRA_ and T_REG_ RNA marker-expressing cells is unlikely to be explained by excessive, cell-type specific cell death (**Sup. Fig. 3A-B**). Observed transcriptional changes correlated with reduced expression of multiple T-cell marker on the cell surface following Panobinostat, but not Vorinostat, treatment, including CD3, CD4, CD45RA (marker for naïve T-cells), CD62L (SELL), CD45RO (marker for memory T-cells), HLA-DR (late marker for activated T-cells), CXCR4, and most pronounced, the cytokine receptor IL7R/CD127 (**Sup. Fig. 3C**). CCR5 and CD69 cell surface expression were mildly increased in the context of Panobinostat treatment, corroborating previous reports of transient CD69 expression following Panobinostat treatment from *in vivo* (44) and *ex vivo* (38) studies (**Sup. Fig. 3D**). Together, among the two HDACi treatments conducted under indicated experimental conditions, Panobinostat induced a massive rearrangement of CD4^+^ T-cell subset marker expression, both on the transcriptional and translational level.

### Panobinostat treatment induces broad transcriptional down-modulation of genes involved in T-cell signalling and a subset of genes implicated in innate immunity

Vorinostat treatment induced modest gene expression changes, with expression of 1.388 genes induced and 312 genes decreased (**Fig. 2A**). In contrast, Panobinostat treatment altered the expression of a total of 10.119 genes. Comparing the two HDACi treatments, 4.078 genes were up-regulated and 5.124 were down-regulated in Panobinostat-compared to Vorinostat-treated cells. Focussing on genes with >2-fold change in expression, cultures treated with individual HDACis displayed a substantial overlap of DEGs, suggesting that transcriptomic alterations were partially similar (**Fig. 2B**). Gene sets previously associated with Notch signalling, synaptic vesicle trafficking and Alzheimer’s disease-presenilin were enriched in Panobinostat-treated cell cultures (**Fig. 2C**). Pathways that were downregulated in the context of Panobinostat treatment were associated with T-cell activation, cytokine- and chemokine-mediated inflammation and JAK/STAT signalling (**Fig. 2C**), in line with reduced activity of multiple transcription factors in all T-cell subsets essential for mediating innate and T-cell-specific immune responses, such as IRF9, STAT1, STAT2, NFKB1 and STAT5, GATA3, RELA, respectively (**Fig. 2D**). Interestingly, activity scores of IRF3 and STAT6 were increased in T_N_- and T_CM_-cell subsets (**Fig. 2D**), suggesting partial, subset-specific activation of innate signalling cascades. Vorinostat failed to enrich or deplete specific gene sets, corroborating the minor impact of Vorinostat at the applied concentration on the overall T-cell transcriptomic profile.

**Figure 2.**
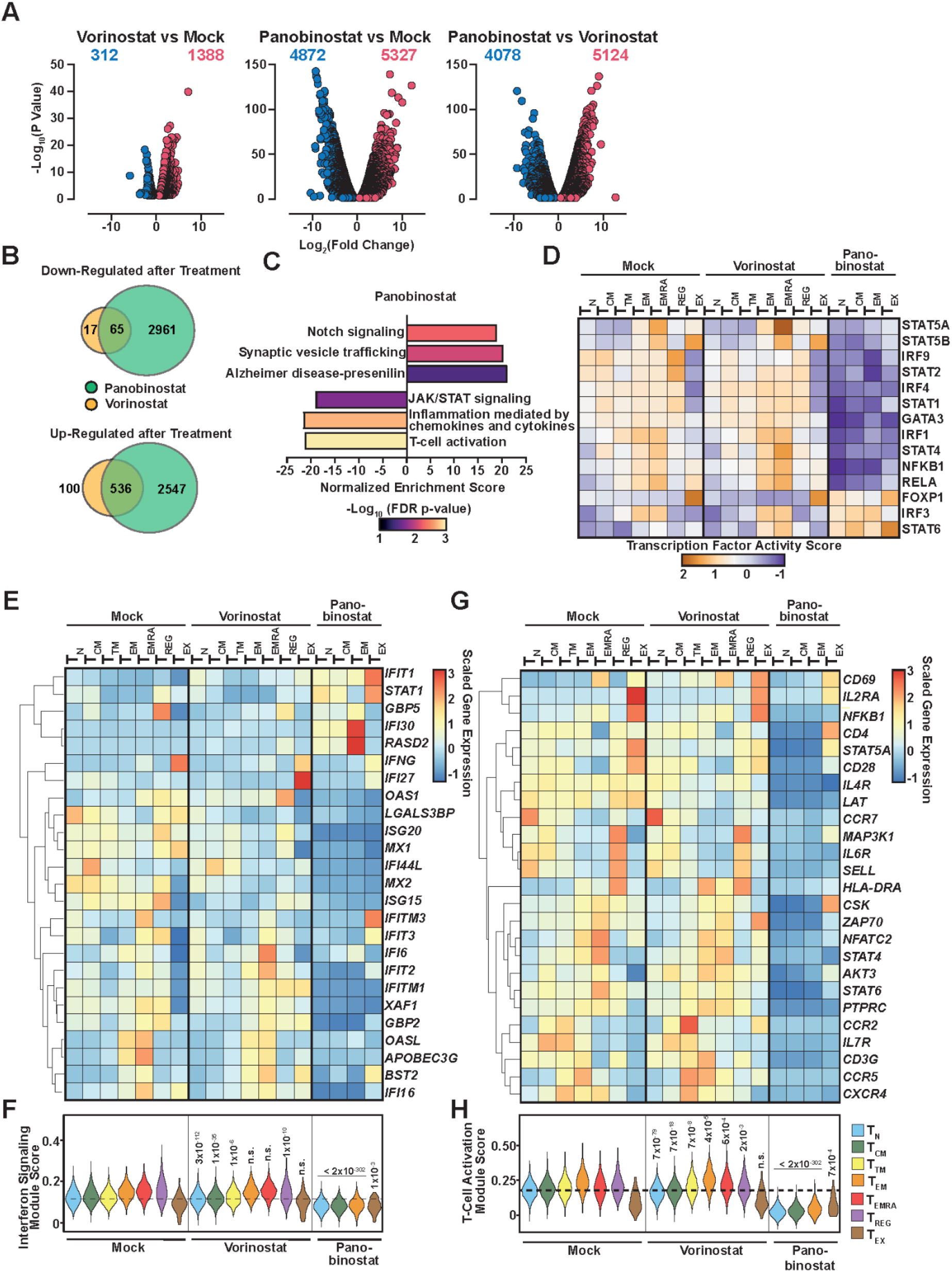
Panobinostat treatment induces broad transcriptional down-modulation of genes involved in T-cell signalling and a subset of genes implicated in innate immunity. **(A)** Volcano plots showing DEGs with a p-value < 0.05. **(B)** Overlap of genes significantly up- or down-regulated (p-value < 0.05, fold change > 2) in Vorinostat and Panobinostat-treated samples compared to mock treatment. **(C)** Pathway analysis of DEGs (p-value < 0.05) after Panobinostat treatment. Shown is the Normalised Enrichment Ration (NER), representing the degree of overrepresentation of a given gene set in the DEG list. Colours indicate the -Log_10_(FDR p-value). **(D)** Transcription factor activity analysis of selected transcription factors in the indicated T-cell subsets. **(E)** Heatmap showing scaled average expression of selected ISGs in the indicated T-cell subsets. Genes are ordered based on correlation, clustering is indicated with lines on the left. **(F)** Average IFN signalling module score in the indicated T-cell subsets. Dashed line shows the average module score of mock-treated T_N_-cells. Numbers indicate the p-value calculated using Wilcoxon signed-rank testing for each subset from the LRA-treated samples compared to the same subset of the mock-treated samples. **(G)** Heatmap showing scaled average expression of selected genes required for T-cell-mediated immunity in the indicated T-cell subsets. Genes are ordered based on correlation, clustering is indicated with lines on the left. **(H)** Average T-cell activation module score in the indicated T-cell subsets. Dashed line shows the average module score of mock-treated T_N_-cells. Numbers indicate the p-value calculated using Wilcoxon signed-rank testing for each subset from the LRA-treated samples compared to the same subset of the mock-treated samples.

Base-line expression of IFN-stimulated genes (ISGs) in untreated cells was to a certain extent subset-specific (**Fig. 2E**). For example, *LGALS3BP* and *IFI44L* displayed the highest expression in resting T_N_- and T_CM_-cell subsets, respectively, while *OASL* and *APOBEC3G* expression was highest in T_EM_- and T_EMRA_-cells. The observations on the individual ISG expression level were corroborated by the IFN Module Score that takes expression of multiple genes associated with IFN signalling into account. T_N_-, T_CM_- and T_TM_-cells displayed an overall modest, T_EM_-, T_EMRA_- and T_REG_-cells gradually increasing and T_EX_-cells a relatively low IFN Module Score in mock-treated CD4^+^ T-cells (**Fig. 2F**). Overall, Vorinostat treatment only mildly affected the ISG expression pattern and IFN Module Score (**Fig. 2E-F**). In contrast, Panobinostat-exposed CD4^+^ T-cells displayed drastically decreased levels of the majority of ISGs, including *ISG20*, *MX1*, *IFI44L*, *IFITM1*, *XAF1*, *GBP2*, *IFI16* throughout all subsets and reduced expression of *IFI27*, *IFITM3*, *IFIT3*, *IFIT2* and *BST2* in T_N_-, T_CM_- and T_EM_-cells, with an opposite trend in T_EX_-cells (**Fig. 2E**). However, *IFIT1, STAT1, IFI30 and RASD2* expression was increased after Panobinostat treatment in all subsets, with *IFIT1* and *STAT1* expression found to be most pronounced in T_EX_-cells, and *IFI30* and *RASD2* expression to be particularly induced in T_EM_-cells (**Fig. 2E**). Interestingly, we previously reported expression of these four genes specifically to be upregulated in the context of base-line cGAS activity (45), which drives IRF3-controlled gene expression, suggesting a partial and CD4^+^ T-cell subset-specific activation of IRF3 following Panobinostat treatment. Nevertheless, the overall IFN Module Score taking the average expression of hundreds of ISGs into account was decreased in all Panobinostat-treated T-cell subsets, while it was largely maintained upon Vorinostat treatment (**Fig. 2F**).

We established a drastic down-regulation of genes involved in T-cell signalling and activity following Panobinostat, but not Vorinostat treatment (**Fig. 2G-H**). As expected under mock treatment conditions, expression of genes whose products regulate and mediate T-cell signalling and activation was clearly T-cell subset-specific (**Fig. 2G-H**). Panobinostat treatment induced downregulation of expression of multiple T-cell function-specific genes, including *IL4R*, *LAT*, *IL6R*, *ZAP70*, and those for which we had detected reduced protein expression on cell surface (CD62L (*SELL*), HLA-DRA, CD45RA/CD45RO (*PTPRC*), IL7R, CCR5 and CD3 (*CD3G*)). Some T-cell-specific genes were specifically upregulated in T_EX_-cells, including *CD4*, *MAP3K1*, *CSK* and *AKT3*. In accordance with the transient nature of CD69 expression upon Panobinostat treatment (**Sup. Fig. 3D**), *CD69* mRNA expression at 48 hours equaled those of mock-treated cells. The overall trend of reduced T-cell activation-specific gene expression was reflected in a T-cell Activation Module Score of Panobinostat-treated cells that was reduced to below the average of mock- and Vorinostat-treated cells (**Fig. 2H**).

In order to test to which extent HDACi-imposed gene expression changes can be recapitulated in a more amenable HIV-1 latency cell system, we treated J1.1 T-cells, that harbour at least two replication-competent HIV-1 proviruses per cell (46,47) with Vorinostat and Panobinostat (**Sup. Fig. 4**). Here, we applied concentrations of HDACis (16 μM Vorinostat and 200 nM Panobinostat) that induced a similar degree of HIV-1 reactivation as judged by intracellular HIV-1 p24 expression (**Sup. Fig. 4A**). Both treatments resulted in a similar number of down- and upregulated genes (**Sup. Fig. 4B**) with a substantial overlap of modulated genes (**Sup. Fig. 4C**). The concentration of applied Panobinostat in J1.1 T-cells (200 nM) approached the one used in primary CD4^+^ T-cells (50 nM) and induced partially similar gene expression changes. Specifically, downregulated activity scores of transcription factors required for T-cell signalling (**Sup. Fig. 4D**), in line with reduced expression of genes related to T-cell activity, such as *CD3G*, *SELL*, *LAT*, *STAT5A* or *ZAP70* and a significantly reduced T-cell Activation Module Score (**Sup. Fig. 4E**). The ISG expression profile following Panobinostat was less well conserved between J1.1 T-cells and CD4+ T-cells. While Panobinostat-treated primary CD4^+^ and J1.1 T-cells shared expression profiles of some ISGs (downregulation of *MX1*, *MX*2, *OAS1*, *XAF1*, *BST2*; upregulation of *GBP5*, *STAT1*, *IFIT1*), they differed regarding expression of *LGALS3BP*, *GBP2*, *IFI44L*, *OASL* (**Sup. Fig. 4F**), resulting in an overall elevated IFN Module Score in J1.1 T-cells as opposed to primary CD4^+^ T-cells.

Conclusively, under these experimental conditions, Panobinostat, and, if applied at higher concentration, also Vorinostat, significantly reduced expression of several genes that are essential for T-cell-specific immune responses and a part of IFN-modulated genes.

### HDAC inhibition imposes a block to CD4^+^ T-cell activation and type I IFN signalling

We hypothesised that HDACi-induced modulation of gene expression is of functional relevance for HIV-1 reactivation and immune recognition of reactivating cells. To investigate the result of HDACi treatment on T-cell activation, we pre-incubated CD4^+^ T-cells from HIV-1-negative donors with HDACis prior to inducing T-cell activation by IL-2/PHA (**Fig. 3A**). Pre-incubation with DMSO or Vorinostat resulted in marked IL-2/PHA-triggered enhancement of the percentage of cells expressing the activation markers CD69, CD25, HLA-DR and also the exhaustion markers TIM-3 and PD-1, indicating successful T-cell activation. Conversely, Panobinostat pretreatment resulted in impaired induction of CD69 expression (0.49-fold changed), and in a complete lack of IL-2/PHA-mediated induction of expression of CD25, HLA-DR, TIM-3 and PD-1 (**Fig. 3A**). To study TCR-induced T-cell activation, we preincubated CD4^+^ T-cells with HDACi followed by treatment with a universal peptide pool in combination with an anti-CD28 antibody. Expression of the early activation marker CD69 after TCR stimulation was slightly, but statistically significantly induced in Vorinostat- (1.65-fold) and to a higher extent in Panobinostat-pretreated samples (5.5-fold), suggesting efficient initiation of T-cell activation in the presence of both HDACis (**Fig. 3B**). Expression of the middle activation marker CD25, the exhaustion marker PD-1 and the late activation marker HLA-DRA increased upon TCR stimulation in the context of Vorinostat pretreatment (2.07-, 2.07- and 1.39-fold change, respectively), suggesting that weak HDAC inhibition does not prevent TCR-specific T-cell activation. However, expression of CD25 and HLA-DRA was severely reduced in Panobinostat-pretreated cultures (0.05- and 0.47-fold change, respectively), suggestive of abortive peptide-induced T-cell activation.

**Figure 3.**
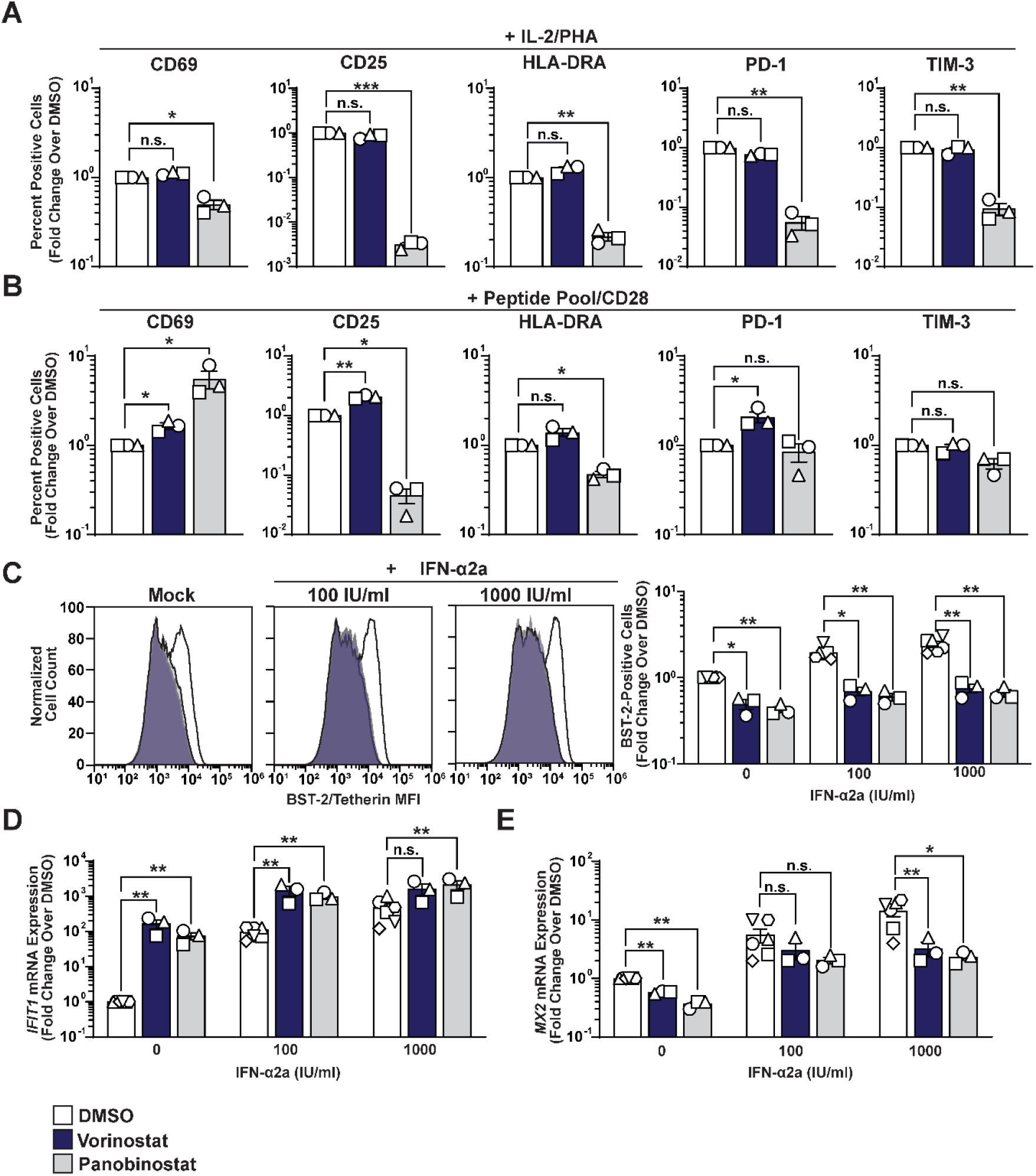
HDAC inhibition imposes a block to CD4^+^ T-cell activation and type I IFN signalling. CD4+ T-cells from uninfected donors were either pretreated with 0.5% DMSO, 500 nM Vorinostat or 50 nM Panobinostat for 16 hours, followed by: **(A)** Stimulation with a peptide pool and anti-CD28 antibody for 48 hours **(B)** Stimulation with IL-2/PHA for 48 hours followed by analysis of CD69, CD25, HLA-DR, TIM-3 and PD-1 cell surface expression by flow cytometry. Shown are data from cells from three donors. CD4^+^ T-cells from uninfected donors were treated with 0.5% DMSO, 8000 nM Vorinostat or 50 nM Panobinostat in combination with increasing concentrations of IFN-α2a for 48 hours before analysing: **(C)** BST2 cell surface levels by flow cytometry. Shown are representative histograms from one experiment (left panel) and quantification from experiments with cells from three donors. **(D)** *IFIT1* mRNA expression by RT-Q-PCR. Shown are data from cells from 3-6 donors. **(E)** *MX2* mRNA expression by RT-Q-PCR. Shown are data from cells from 3-6 donors.

We next examined the impact of HDAC inhibition on type I IFN signalling and ISG expression. Specifically, we treated CD4^+^ T-cells isolated from HIV-1-negative donors with DMSO, Vorinostat or Panobinostat in combination with increasing concentrations of IFN-α2a and assessed cell surface expression changes of BST-2 and changes of *IFIT1* and *MX2* gene expression (**Fig. 3C-D**). We applied an increased concentration of Vorinostat in these experiments to adjust for the inferior potency of Vorinostat compared to Panobinostat that has been reported in previous studies (38,48). Both Vorinostat and Panobinostat significantly dampened base-line BST-2 cell surface expression levels, and both tested concentrations (100 and 1000 IU/ml) of IFN-α2a failed to rescue the HDACi-imposed block to BST2 surface expression (**Fig. 3C**). Interestingly, mRNA expression levels of the two ISGs *IFIT1* and *MX2* showed a divergent phenotype. Individual treatment with Vorinostat and Panobinostat, as well as addition of IFN-α2a, induced and further amplified *IFIT1* mRNA expression, respectively (**Fig. 3D**). In contrast, *MX2* mRNA expression decreased in the context of HDACi treatment, both in the absence and presence of 1000 IU/ml IFN-α2a (**Fig. 3E**). These results were paralleled in latently HIV-1-infected J1.1 T-cells in which HDAC inhibition, but not treatment with the non-HDACi LRA Bryostatin, reverted prior IFN-α2a-induced expression of MX2 in a dose-dependent manner and abolished the negative impact of IFN on early HIV-1 reactivation as judged by intracellular p24 expression (**Sup. Fig. 5A).** Co-administration of increasing doses of IFN-α2a up to 10,000 IU/ml was insufficient to counteract the HDACi-suppressed MX2 expression and did not limit the efficiency of HIV-1 reactivation, in contrast in the context of Bryostatin treatment, IFN treatment induced MX2 expression efficiently and inhibited HIV-1 reactivation in a dose-dependent fashion (**Sup. Fig. 5B**).

Together, HDAC inhibition renders CD4^+^ T-cells largely refractory, and/or less sensitive, to mitogen and TCR stimulation and dampens the base-line and IFN-induced expression of multiple ISGs, including the two prototypic ISGs BST-2 and MX2, while IRF3-driven expression of a small subset of ISGs remains intact and is even exacerbated. We hypothesise that the HDACi-induced cellular phenotypes both at the level of CD4^+^ T-cells as HIV-1 reactivating target cells, and CD4^+^ T-cell effector functions is likely to contribute to the lack of HIV-1-positive T-cell elimination *in vivo* both at the level of CD4^+^ T-cells as HIV-1 reactivating target cells, and CD4^+^ T-cell effector functions.

### Addition of IFN-α2a to Panobinostat results in a tetherin-independent pre-budding defect

The previous data highlighted that HDAC inhibition significantly diminishes IFN-induced signalling, potentially nullifying the negative effect of IFN on HIV-1 replication. To test in detail the impact of IFN on HIV-1 reactivation and multi-round spread upon LRA treatment, J.1. T-cells were primed with a single dose of 50 nM Panobinostat in the presence or absence of IFN-α2a. The percentage of HIV-1 p24 capsid-expressing cells was initially similar, but differed starting from day three on (**Fig. 4A**, **Sup. Fig. 6**). Addition of IFN to Panobinostat reduced the abundance of p24 capsid (**Fig. 4B**) and infectivity (**Fig. 4C**) released into cell culture supernatant, effects detectable from day two and one post-treatment start on, respectively. Two days post HIV-1 reactivation, a time point at which cell-associated p24 capsid expression is statistically indistinguishable (**Fig. 4A**), the morphology of budding sites was similar and intact in both conditions (**Fig. 4D**), however, the incidence of detectable budding events per cell was reduced 2.2-fold in the context of the co-treatment (**Fig. 4E**). Importantly, this observation was obtained in the absence of tetherin-imposed attachment of virions to the cell surface or to each other (**Fig. 4D)**. This tetherin-atypical microscopic appearance, and the lack of IFN-induced tetherin upregulation when Panobinostat is present (**Fig. 4F, G**) strongly argues against tetherin driving this antiviral phenotype, and rather suggests a defect prior to release or a tetherin-independent release defect.

**Figure 4.**
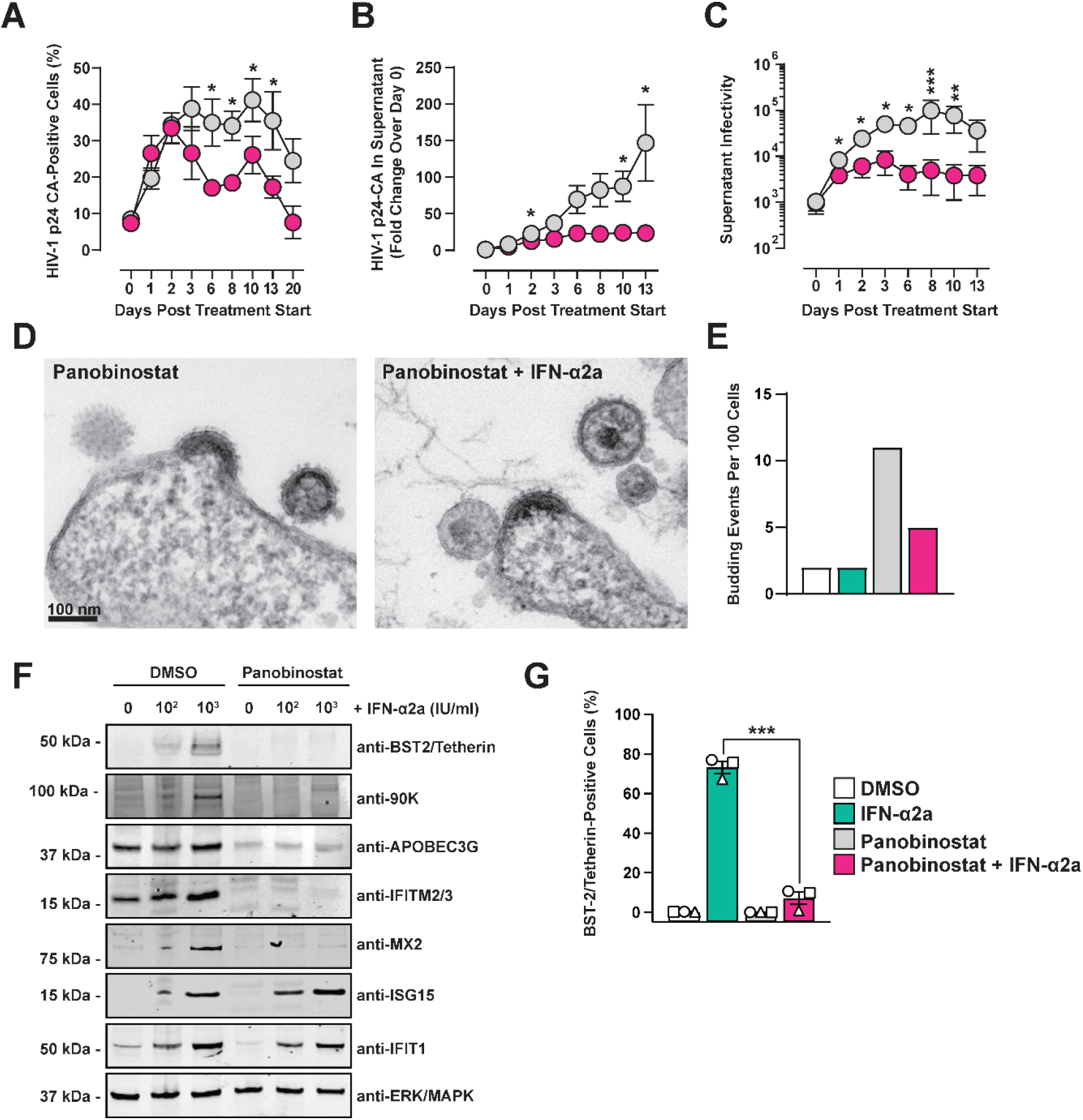
Addition of IFN-α2a to Panobinostat results in a tetherin-independent pre-budding defect. J1.1 T-cells received a single dose of 50 nM Panobinostat alone or in combination with 100 IU/ml IFN-α2a. At indicated time points, samples were analysed for: **(A)** Cell-associated HIV-1 p24 capsid expression by flow cytometry **(B)** HIV-1 p24 capsid abundance in cell culture supernatant by immunoblotting. **(C)** Infectivity in cell culture supernatant using the Tzm-bl reporter cell assay. **(D)** Representative electron micrographs of reactivated J1.1 T-cells 48 hours post reactivation. Scale bar indicates 100 nm. **(E)** Electron microscopy-based quantification of budding events per 100 cells analysed. **(F)** Immunoblot analysis of selected ISG expression in J1.1 T-cells following 48 hours of indicated treatment. **(G)** J1.1 T-cell surface BST-2/Tetherin expression 48 hours following treatment start.

### IFN-induced virus pre-budding block is accompanied by accumulation of HIV-1 Env on cellular and viral membranes

Next, we assessed the nature of the IFN-imposed block prior to HIV-1 budding. Despite increasing concentrations of IFN-α2a, Panobinostat-induced cell-associated HIV-1 p24 capsid expression remained intact (**Fig. 5A**). In contrast, the p24 capsid signal intensity in corresponding supernatants (**Fig. 5B**) and normalised release (**Fig. 5C**) dose-dependently decreased in cultures co-treated with IFN-α2a. Interestingly, combined Panobinostat/IFN-α2a treatment resulted in a higher ratio of gp120 per p24 (**Fig. 5D**), suggesting a higher rate of Env incorporation in released virions in the absence of evidence for an Env processing alteration in corresponding cell lysates (**Sup. Fig. 7A**). Panobinostat, alone or in combination with IFN-α2a, increased the percentage of cells expressing HIV-1 Env on their surface (**Fig. 5E**), in line with increase of cell-associated p24 expression by both conditions (**Fig. 5A**). However, IFN-α2a addition to Panobinostat had a 1.32-fold, statistically significant increase of the mean fluorescence intensity (MFI) of Env signal on Env-positive cells as a consequence, indicating accumulation of HIV-1 Env molecules on the cell surface (**Fig. 2F**). Treatment with the JAK/STAT inhibitor Ruxolitinib preserved the increase of percentage of Env-positive cells (**Sup. Fig. 7B**), but nullified IFN-induced Env accumulation per cell (**Sup. Fig. 7C**), consistent with upregulation of Env requiring functional IFN signalling. Conclusively, combined treatment of Panobinostat and IFN-α2a results in increased quantities of HIV-1 Env on reactivating T-cellś surfaces and released virions.

**Figure 5.**
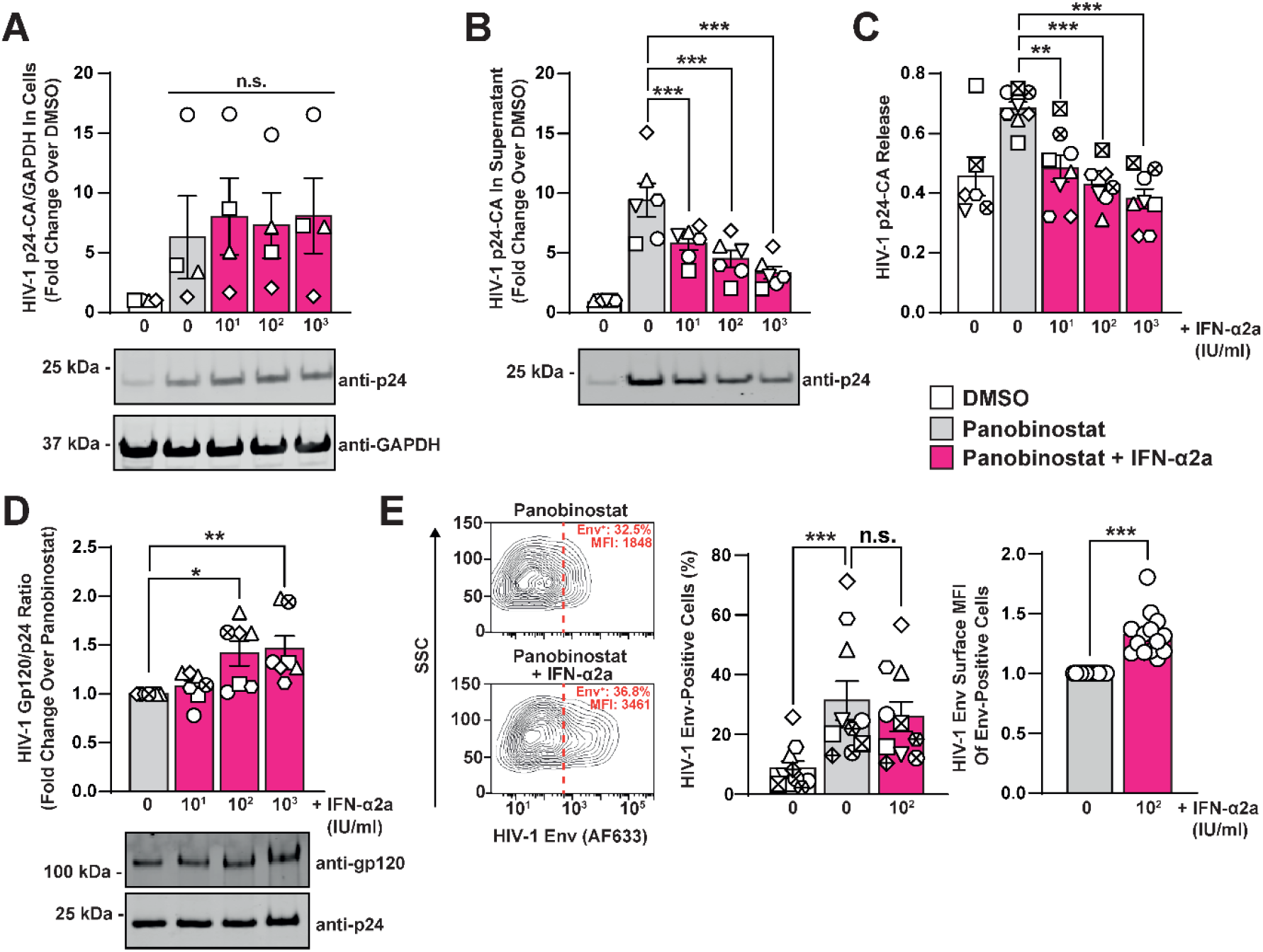
IFN-induced virus pre-budding block is accompanied by accumulation of HIV-1 Env on cellular and viral membranes. J1.1 T-cells were treated with 50 nM Panobinostat in the presence or absence of indicated concentrations of IFN-α2a for 48 hours. Cells and corresponding supernatants were analysed for: **(A)** Relative expression of cell-associated HIV-1 p24-CA by immunoblotting, normalised to cellular GAPDH. **(B)** Relative abundance of extracellular HIV-1 p24-CA by immunoblotting, normalised to corresponding DMSO controls. **(C)** Virus release was quantified by normalising extracellular HIV-1 p24-CA signal intensity to the sum of cell-associated and secreted HIV-1 p24-CA. **(D)** HIV-1 gp120 signal intensities from cell culture supernatants were quantified by immunoblotting and normalised to corresponding HIV-1 p24-CA levels. **(E)** Representative dot plots of intact J1.1 T-cells immunostained for surface HIV-1 Env with indicated percentage of Env-positive cells and mean fluorescence intensity (MFI) of Env-positive cells. Quantification of HIV-1 Env-positive cells (middle) and the fold change of Env MFI in Env-positive cells over Panobinostat-treated cells (right) are shown for multiple experimental replicates. Experiments were conducted in 5-13 independent replicates.

### Addition of IFN-α2a to Panobinostat enhances the susceptibility of reactivating T-cells to NK-mediated antibody-dependent cellular cytotoxicity

Being the sole viral antigen presented on the virus-producing cell surface, HIV-1 Env protein serves as the major target of antibody-mediated responses. We therefore examined whether the increased surface expression of HIV-1 Env in the context of IFN-α2a treatment renders cells more susceptible to ADCC. Specifically, we co-cultured HIV-1-reactivating, cell tracker^TM^ Green-CMFDA-stained J1.1 T-cells and freshly isolated PBMCs in the presence of serum from PLHIV, and loss of HIV-1 p24 capsid-positive cells was monitored (**Sup. Fig. 8A**). ADCC decreased with increasing dilutions of the anti HIV-1-Env antibody-containing serum, independent of the reactivation regimen applied (**Fig. 6A-C**). Interestingly, Panobinostat treatment failed to significantly enhance ADCC as compared to DMSO treatment, suggesting that it increases the quantities of HIV-1 p24- and Env-positive cells, but not their relative susceptibility to antibody-mediated elimination. In contrast, J1.1 T-cells reactivated in the presence of IFN-α2a scored overall higher ADCC values. T-cells co-cultured with isolated NK cells, but not monocytes, displayed a statistically significant increase of ADCC in the context of IFN-α2a co-treatment as compared to Panobinostat only-treatment (**Sup. Fig. 9A, B**), arguing for a predominant role of NK cells rather than monocytes in mediating T-cell killing. In line with this, CD107a surface exposure and CD16 downregulation, which strongly correlates with NK cell activation, cytokine production and target cell lysis (47–49), were highly induced in the NK cell population of PBMCs co-cultured with J1.1 T-cells (**Sup. Fig. 9C-E**). Co-treatment of J1.1 T-cells with Panobinostat and IFN-α2a, however, did not result in notable differences of the two Panobinostat-treated conditions in regards to NK cell effector activation despite increased target cell killing in the co-culture (**Sup. Fig. 9D-E**), arguing for IFN-a2 increasing the sensitivity of target cells to killing rather than modulating effector cell activity.

**Figure 6.**
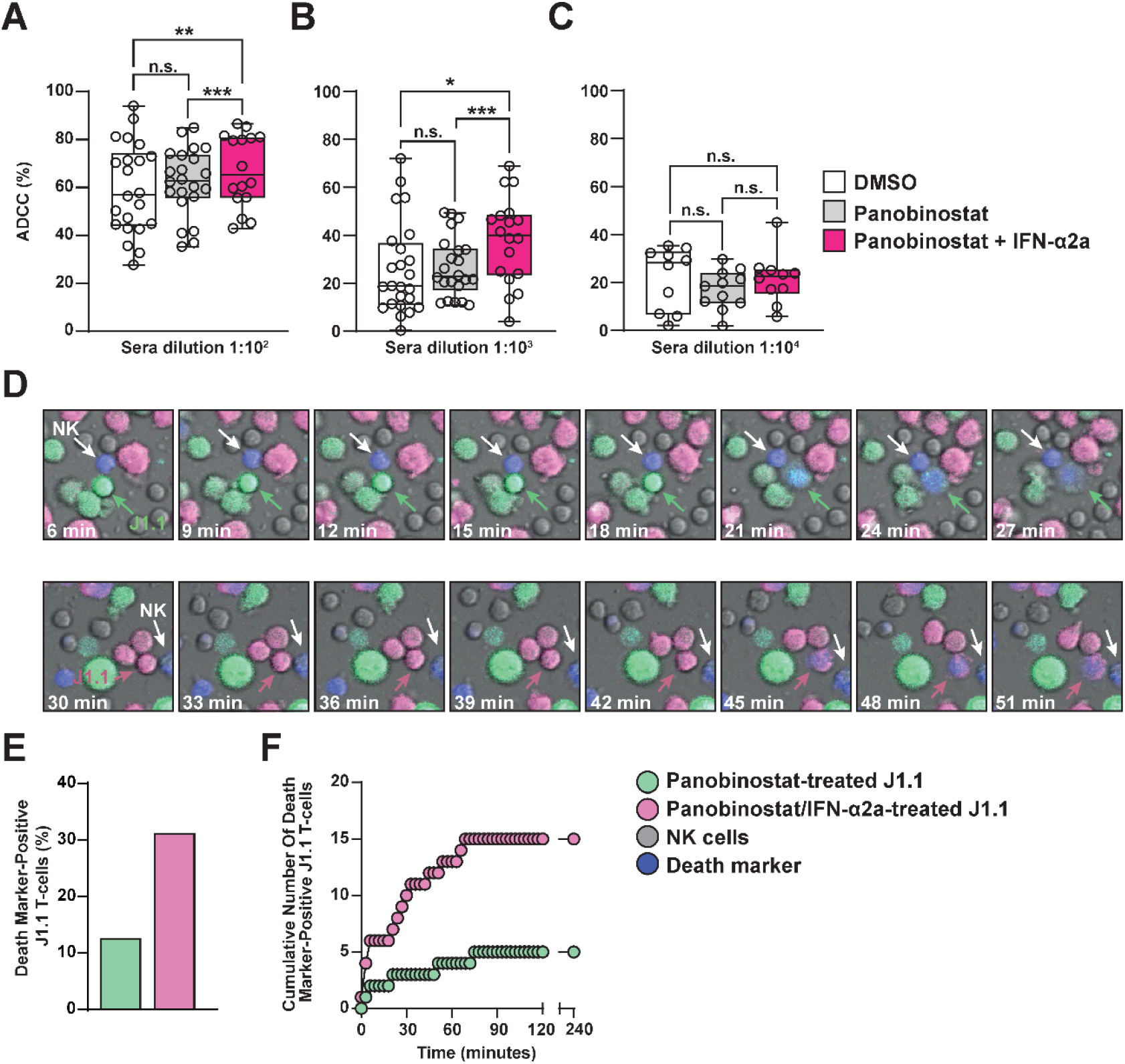
Addition of IFN-α2a to Panobinostat enhances the susceptibility of HIV-1-reactivating cells to antibody-dependent cellular cytotoxicity via NK cells. (**A-C**) J1.1 T-cells were treated with DMSO, 50 nM Panobinostat alone or in combination with 100 IU/ml IFN-α2a for 48 hours before culturing them with PBMCs in the presence of indicated dilutions of sera from PLHIV. Following four hours of co-culture, cells were immunostained for cell-associated HIV-1 p24 CA expression and percentage of ADCC was calculated based on the quantitative difference of HIV-1-p24-positive cells in the identical co-culture merely differing in the presence or absence of serum. Boxes extend from the 25th to 75th percentiles of 10-24 repetitions, the lines indicate the median, error bars represent the minimum and maximum values obtained and symbols the individual experimental replicates. (**D**) - (**F**) J1.1 T-cells were treated with 50 nM Panobinostat alone or in combination with 100 IU/ml IFN-α2a for 48 hours and individually stained with CellTracker^TM^ Green-CFMDA and Deep Red, respectively. Stained J1.1 T-cells were co-cultured with non-stained, freshly isolated NK cells at a 1:1:1 ratio in the presence of a cell death marker (blue) and a 1:100 dilution of sera from PLHIV containing anti-HIV-1 Env antibodies. The co-culture was monitored using live-cell imaging for four hours with images acquired every three minutes. (**D**) Representative time series of NK cell/J1.1 T-cell interactions and subsequent killing of J1.1 T-cells. (**E**) Percentage of dead J1.1 T-cells after four hours, normalised to the number of living J1.1 T-cells in the first frame. (**F**) Cumulative number of death marker-positive cells in each J1.1 T-cell population over time.

We next investigated whether IFN-α2a addition modulates the kinetics of NK cell-effected ADCC against Panobinostat-treated J1.1 T-cells using live-cell imaging. Specifically, J1.1 T-cells were treated either with Panobinostat only or Panobinostat in combination with IFN-α2a for 48 hours, followed by differential staining of these two J1.1 T-cell cultures using Cell Tracker™ Green CMFDA and Cell Tracker^TM^ Deep Red, respectively. Cells were then co-cultured with freshly isolated, unstained NK cells at a 1:1:1 ratio in the presence of serum from PLHIV and a cell death marker (blue) and imaged every three minutes for a total of four hours (**Sup. Movie 1**). The starting frame contained similar numbers of both J1.1 T-cell cultures (47 green cells, 51 red cells). As expected, the overall number of cell death marker-positive J1.1 T-cells increased over time, indicating ADCC. We did not detect profound differences regarding the duration of individual NK cell/J1.1 T-cell interactions depending on the T-cell line pretreatment (**Sup. Movie 1**). However, the percentage of cells dying until the end of the experiment, as judged by acquisition of the blue fluorescence and visual disappearance was clearly increased for cells receiving Panobinostat/IFN-α2a co-treatment (31.4%) as compared to cells pre-treated with Panobinostat alone (12.8%) (**Fig. 6E**). The number of Panobinostat/IFN-α2a-treated J1.1 T-cells showing signs of cell death increased rapidly and plateaued at 69 minutes after the start of the imaging (**Fig. 6F**). In contrast, the number of Panobinostat-treated J1.1 T-cells with signs of cell death increased only moderately and plateaued after 75 minutes (**Fig. 6F**). Despite the high mobility and intact shape of the NK cells, no more J1.1 T-cells were eliminated 75 minutes after start of the co-culture, suggesting that the remaining J1.1 T-cells either did not reactivate HIV-1 expression or were otherwise resistant to killing. Of note, a substantial fraction of NK cells that otherwise presented an intact cell shape and high mobility acquired the blue death marker over time (**Sup. Movie 1**). We hypothesise that the processes of degranulation and re-internalization of granules in highly active NK cells allow the uptake of the cell death dye from the medium, not necessarily reflecting cell death. This is in line with the observation that specifically the highly mobile, J1.1 T-cell-interacting NK cells stained positive for the cell death marker, the same population of cells that is thought to have a high cytotoxic potential.

Overall, our observations are in line with a scenario in which addition of IFN-α2a to a Panobinostat-mediated HIV-1 reactivation regimen renders HIV-1-positive cells more prone to NK cell-mediated ADCC, in line with higher levels of the major antibody target Env on the cells’ surface, despite similar levels of HIV-1 reactivation.

### IFN-α2a-enhanced HIV-1 Env cell surface accumulation and ADCC sensitization occur in the context of HDACi-, but not PKC agonist-mediated HIV-1 reactivation

We next tested whether IFN-α2a-mediated upregulation of HIV-1 Env and sensitivity of cells to immunological elimination occurs in conjunction with a specific class of LRA or, on the contrary, universally upon HIV-1 latency reversal. Like Panobinostat treatment, individual treatment of cells with the HDACis Vorinostat and Romidepsin, and the PKC agonist Bryostatin strongly increased the percentage of HIV-1 p24 capsid-positive cells, while the BET inhibitor JQ1 was inferior in reversing proviral quiescence (**Sup. Fig. 10A**). Overall, the percentages of HIV-1 Env-positive cells induced by Vorinostat, Romidepsin and JQ1 treatment followed the percentages of p24-positive cells. Surprisingly Bryostatin treatment, both in the absence or presence of IFN-α2a, failed to increase the percentage of Env-positive cells, despite clear HIV-1 reactivation at the level of p24 capsid (**Sup Fig. 10B**), and accordingly, both treatments did not alter susceptibility to ADCC which, surprisingly, was clearly detectable despite low surface Env levels (**Sup. Fig. 10C**). In summary, our findings suggests that IFN-α2a co-treatment enhances Env cell surface expression in the context of HDACi and the iBET JQ1, but not Bryostatin, pointing towards LRA class-specific differences in HIV-1 Env cell surface targeting and/or regulation.

### The ability to restrict HIV-1 budding and facilitate elimination of HIV-1-positive T-cells is type I IFN protein-specific

Although previous HIV-1 cure strategies focused on incorporation of IFN-α2a or -α2b in shock-and-kill approaches, *in vitro* work clearly demonstrates that other members of the type I IFN family, including IFN-α6, IFN-α14 and IFN-β, display superior anti-HIV-1 properties (49–52). To investigate the breadth of type I IFN proteins to enhance susceptibility of HIV-1-reactivating cells to elimination, we selected IFN-α6, IFN-α14 and IFN-β due to their previously reported high anti-HIV-1 activity (49,50,52), IFN-α16 for its comparatively weaker activity (49,50,52) and IFN-α2 as a reference. IFN-α14 and -β treatment resulted in strongest ISG induction, as judged by quantification of *IFIT1* mRNA and BST-2 protein expression (**Sup. Fig. 11**). Upon co-treatment with Panobinostat, all tested type I IFNs, particularly IFN-α14 and -β, significantly decreased secreted infectivity (**Fig. 7A**). Surprisingly, all IFNs shared the ability to increase the HIV-1 Env surface MFI (**Fig. 7B**), but differed in their ability to sensitise T-cells for elimination, with IFN-α14 and -β treatment resulting in highest ADCC rates (**Fig. 7C**). It is worth mentioning that the activity of the in-house generated IFN subtypes used in Fig. 7 was determined using an ISRE-reporter cell line in contrast to commercial IFN-α2a (Roferon, used in the rest of the study) whose activity is quantified in a cytopathic effect inhibition assay. These differences hindered a direct comparison of the two IFN-α2 potencies and potentially explain the relative lack of IFN-α2 in inducing ADCC under these experimental conditions (**Fig. 7C**). The antiviral activity of individual type I IFN proteins correlated with the induced susceptibility to ADCC. The degree of upregulation of cell surface Env levels did not predict ADCC susceptibility (**Fig. 7D**), suggesting that IFN-mediated enhancement of surface Env levels is required, but not sufficient for sensitising cells to elimination.

**Figure 7.**
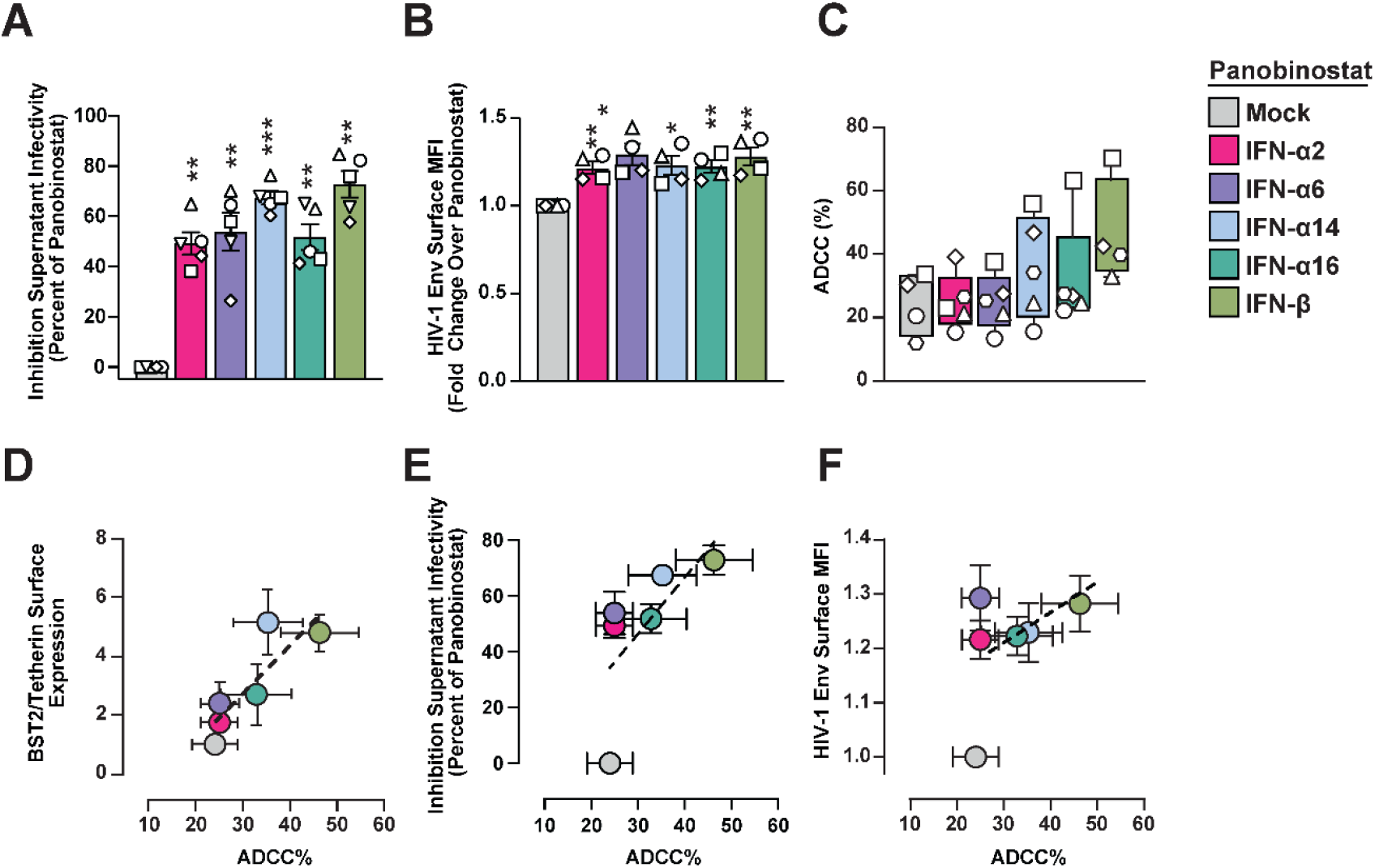
The ability to restrict HIV-1 budding and facilitate elimination of HIV-1-positive T-cells is type I IFN protein-specific. J1.1 T-cells were treated with 50 nM Panobinostat in combination with 100 IU/ml of the indicated IFNs for 48 hours. (**A**) Inhibition of supernatant infectivity was quantified using a Tzm-bl reporter assay, values were normalised to the corresponding Panobinostat-only conditions. (**B**) Fold change of Env MFI in Env-positive cells over Panobinostat-treated cells. (**C**) J1.1 T-cells were co-cultured with PBMCs in the presence of serum from PLHIV (dilution 1:1000) to monitor and quantify ADCC. (**D**) - (**F**) Pearson’s correlation analysis of ADCC % with BST2/Tetherin surface expression, supernatant infectivity and HIV-1 Env surface expression levels.

In summary, we identified a superior ability of the clinically yet underexplored members of the type I IFN family, IFN-α14 and -β, to enhance susceptibility to elimination of HIV-1-reactivating T-cells which correlates with their ability to inhibit HIV-1 budding in the context of Panobinostat reactivation and to accumulate viral Env on the HIV-1-reactivating cells’ surface.

## DISCUSSION

The overarching aim of LRA treatment in the context of HIV-1 cure is reinitiation of HIV-1 mRNA and protein expression to a degree that is sufficient to render a maximum of HIV-1-reactivating cells immunologically visible, making them susceptible to elimination. The ability of HDACis to reverse HIV-1 latency *in vitro* and *in vivo* (12,36–38,53) involves hyperacetylation of histones, which leads to opening of cellular chromatin and subsequently modulation of gene expression, including transcriptional reactivation from dormant HIV-1 genomes (54,55). LRA treatment *per se* influences the expression profile of a plethora of cellular genes, with to date poorly defined consequences on cellular processes affecting HIV-1 RNA stability, HIV-1 protein expression, trafficking and cell surface presentation, and finally susceptibility of HIV-1-reactivating cells to cytotoxicity. An optimal shock-and-kill regimen should, on the one hand, mount a proviral milieu that facilitates post-integration steps of the virus replication cycle. On the other hand, in a second step, an ideal shock-and-kill regimen must render reactivating T-cells susceptible to recognition and killing by CD8^+^ T-cell and NK cell-mediated cytotoxicity, processes that are orchestrated in the effector cell, among others, by functional type I IFN signalling (27,56). Treatment of CD4^+^ T-cells with Panobinostat resulted in a markedly impaired ability to respond to activation stimuli, including TCR/CD28 and IL-2/PHA, in line with reports of HDACi treatment-induced antigen-specific anergy of mouse lymphocytes (57). Given that Panobinostat treatment compromises functionality of NK cells (58) and CD8^+^ T-cells (59,60), it is tempting to speculate that it also affects the efficiency of peptide processing and presentation in HIV-1-reactivating CD4^+^ T-cells by MHC-I molecules. Along this line, HDACi treatment has been proposed to reduce cytosolic peptidase activities in *ex vivo*-HIV-1-infected CD4^+^ T-cells, resulting in modulated antigen presentation to CD8^+^ T-cells (61). Together, we suspect that these effects compromise the overall effector functions of several immune cell types that would otherwise contribute to elimination of HIV-1-reactivating T-cells.

A second striking consequence of Panobinostat was a largely CD4^+^ T-cell subset unspecific, broad, though not entire shut-down of components of IFN-related signalling pathways and functions, which is consistent with the known requirement of functional histone deacetylation for IFN signalling, but not for IRF3-mediated gene expression programmes (62). This leads us to propose to re-consider the contribution of several antiviral genes when analysed in the context of HDACi-induced reactivation, since expression of several ISGs is repressed by HDACi treatment even in the presence of high concentrations of type I IFN. A striking example is our observation of an IFN-induced release defect. Iin the context of Panobinostat treatment, tetherin expression is downregulated in most CD4^+^ T-cell subpopulations except the exhausted CD4+ T-cell subset, fully prevented even when type I IFN is exogenously added, and the microscopic appearance of stalled budding events resembles more those described in the context of ESCRT component defects (63). Together, these data argue rather for a to date unknown ISG product whose IFN dependency is still preserved despite HDAC inhibition.

Type I IFN treatment in conjunction with Panobinostat-mediated HIV-1 reactivation was followed by increase of HIV-1 Env molecule density on viral and HIV-1-reactivating T-cells’ surfaces. HIV-1 tightly regulates the presentation of Env on the cell surface to minimise the vulnerability against humoral responses while maintaining infectivity (64). The observed Env cell surface accumulation may be an indirect consequence of the pre-budding defect, where fewer virions are detached from the producer cell, resulting in excess HIV-1 Env at the plasma membrane, and/or may be a direct consequence of the HDACi/IFN treatment combination on HIV-1 Env trafficking and recycling. The rate of internalisation of Env molecules and their conformation presented at the cell surface have been reported as determinants of the HIV-1-infected cell’s susceptibility to ADCC (65–67). Our data points towards a scenario in which, in the context of Panobinostat treatment, increased Env expression on reactivating T-cells, likely with contribution of a potential ISG product, sensitises HIV-1-reactivating T-cells for NK cell-mediated elimination. This would represent a novel mechanism by which type I IFN sensitises, rather than protects, target cells for elimination. On the contrary, several studies reported type I IFNs to protect T-cells from NK cell-mediated cytotoxicity by processes involving upregulation of expression of selected inhibitory NK-cell-receptor ligands (68) and reduction of expression of natural cytotoxicity triggering receptor 1 (NCR1) ligands as evasion mechanism (69). However, these studies were not conducted in the context of pharmacological reactivation of HIV-1. Specifically, a higher proportion of Panobinostat-only treated cells remained viable at the end of the co-culture as compared to IFN-co-treated cells, indicating an intrinsic resistance to ADCC-mediated killing that was partially overcome by the addition of IFN. Importantly, we did not observe differences in the degranulation activity or cellular activation of NK cells retrieved from co-cultures with either Panobinostat- or Panobinostat/IFN-treated J1.1 cells, suggesting that the impact of IFN on NK cell effector activity is neglectable under these experimental conditions. To date, only tetherin has been identified as an IFN-stimulated factor that augments ADCC responses against HIV-1-infected cells (67,70) however, it is an unlikely candidate in the context of Panobinostat treatment due to its virtually absent expression in the context of HDAC inhibition. Adding another layer of complexity, susceptibility to immune-mediated clearance is not determined be mere quantities of surface-accessible Env, as evidenced by our data showing clear antibody-dependent killing despite low cell surface levels of Env on target T-cells following Bryostatin-mediated latency reversal, and IFN-protein-specific ADCC susceptibilities despite similar Env cell surface levels following Panobinostat treatment. The characterisation of cellular and viral factors that determine the susceptibility of HIV-1-reactivating cells to ADCC and how these factors are manipulated by individual pharmacologic latency reversal strategies are key questions to be addressed in future studies.

In the context of HIV-1 shock-and-kill cure approaches, HDACis belong to the most thoroughly studied class of LRAs and are well tolerated *in vivo* (71,72), though administration of HDACis appears to inhibit essential immune effector functions of CD8^+^ T-cells (60), NK cells (58,73) and reduce susceptibility of target T-cells to cytotoxic elimination as shown in our study. However, the overall high potency of HDACis to reactivate HIV-1 and the lack of alternative LRAs with better *in vivo* performance encourages the continued use of HDACis in newly launched clinical trials, especially in combination with immunomodulatory drugs to counterbalance the HDACi-mediated negative effects. Currently, HDACis are being tested in combination with pegylated IFN-α2a (Clinical trial NCT02471430), broadly neutralising antibodies (Clinical trial NCT02850016) (74,75) and therapeutic vaccine candidates (Clinical trials NCT02092116 and NCT02616874) (76). Within this conceptually broad field, our study is the first proof-of-concept that provides a mechanistic rationale that justifies incorporating type I IFNs in shock-and-kill settings to enhance the susceptibility of HIV-1-reactivating T-cells to immune-mediated elimination. Importantly, the superior ability of the clinically underexplored IFN-α14 and IFN-β over IFN-α2a to sensitise T-cells for elimination justifies their advancement for clinical trials. In conclusion, combining potent LRAs with immunomodulatory molecules, such as type I IFNs, presents a promising strategy to overcome the intrinsic and/or virus-mediated resistance of HIV-1-reactivating T-cells to NK cell-mediated killing which would be an instrumental step towards effective shock-and-kill cure strategies.

## MATERIAL AND METHODS

### Cell lines and primary cells

HEK293T and Tzm-bl cells were obtained from ATCC and the NIH AIDS Reagents Program and maintained in Dulbecco’s modified Eagle’s medium (DMEM) supplemented with 10% heat-inactivated fetal calf serum (FCS), 100 IU/ml Penicillin/Streptomycin and 2 mM L-Glutamine in a 5% CO_2_ atmosphere at 37°C. Jurkat and J1.1 T-cells were obtained from the NIH AIDS Reagents Program and cultivated in RPMI 1640 supplemented with 10% heat-inactivated FCS, 100 IU/ml Penicillin/Streptomycin, 2 mM L-Glutamine, 1x MEM non-essential amino acids and 1 mM sodium pyruvate in a 5% CO_2_ atmosphere at 37 °C.

Withdrawal of blood samples from healthy human donors and cell isolation were conducted with approval of the local ethics committee (Ethical review committee of Charité Berlin, vote EA4/167/19). Withdrawal of blood samples from aviremic PLHIV and cell isolation were conducted with approval of the local ethics committee (Ethical review committee of Charité Berlin, votes EA2/105/05 and EA2/024/21) in the context of the HIV-1 Seroconverter Study of the Robert-Koch-Institute (77). Available clinical information is indicated in Supplementary Table 1. Human PBMCs, CD4^+^ T-cells, NK cells and monocytes were isolated from EDTA-anticoagulated blood by Ficoll-Hypaque centrifugation or using the EasySep Direct Human CD4^+^ T-cell, Human NK cell or Human Monocyte CD14^+^ Isolation kits (STEMCELL Technologies), respectively. Primary cells were cultured at 10^6^ cells/ml in RPMI 1640 containing 10% heat-inactivated FCS, 100 IU/ml penicillin/streptomycin, 2 mM L-Glutamine, 1% MEM non-essential amino acids and 1 mM sodium pyruvate.

### Reagents and inhibitors

The following commercial reagents were used throughout this study: Bryostatin I (Sigma Aldrich), CEFX Ultra SuperStim Pool (#PM-CEFX-1, JPT), IFN-α2a (Roche), IFN-β (Roche), IL-2 (Merck), JQ-1 (Sigma Aldrich), Panobinostat (Cayman Chemical), PHA (Thermo Scientific), Romidepsin (Selleckchem), Ruxolitinib (STEMCELL Technologies) Vorinostat/SAHA (Abcam). Efavirenz was obtained from the NIH AIDS Reagent Program. Recombinant IFN-α subtypes 2, 6, 14 and 16 were expressed in *Escherichia coli*, purified by anion-exchange and size exclusion chromatography and tested for absence of endotoxins (50). IFN units were determined using a human retinal pigment epithelial reporter cell line stably expressing a plasmid containing the IFN-stimulated response element driving a luciferase reporter gene.

### Treatment of CD4^+^ T-cells for single cell RNA-sequencing

CD4^+^ T-cells were mock-treated (0.05% DMSO) or treated with Vorinostat (500 nM), Panobinostat (50 nM) or IL-2 (20 IU/ml) and PHA (1 μg/ml) for 48 hours before subjecting them to the single cell RNA-sequencing pipeline. J1.1 T-cells were treated with 1.6% DMSO (DMSO I), 16000 nM Vorinostat, 0.2% DMSO (DMSO II) or 200 nM Panobinostat for 40 hours before subjecting them to single cell RNA-sequencing.

### Single cell RNA-sequencing

Single cell RNA-Seq libraries were prepared with the 10x Genomics platform using the Chromium Next GEM Single Cell 3’ Reagent Kits v.3.1 following manufacturer’s instructions. Quality control of the libraries were performed with the KAPA Library Quantification Kit and Agilent TapeStation. Libraries were sequenced on a HiSeq4000 using the following sequencing mode: read 1: 28 bp, read 2: 91-100 bp, Index i7: 8 bp. The libraries were sequenced to reach ∼20 000 reads per cell.

### Single cell RNA-sequencing data analysis

FASTQ files from the sequencing protocol were processed using the Cell Ranger pipeline v 3.1.0 (10X Genomics) and further analysed using the Seurat v3.1.4 package (78) in R v3.6 (R Core Team, 2017). Reads from cells isolated from PLHIV were aligned to the human genome (GRCh38), while reads from J1.1 T-cells were aligned to a custom reference consisting of the genomic HIV-1 RNA (LAV-1, GenBank: K02013.1) appended to GRCh38 to allow for capture of viral RNAs. The data was processed using the SCTransform workflow as outlined by the Seurat developers. Cells were clustered using Louvain clustering in a UMAP projection. For the CD4^+^ T-cells from PLHIV, T-cell subsets were identified based on marker gene expression: T_N_ cells (CD3D^+^, CD8A^-^, CCR7^+^, S100A4^low^) (39), T_CM_ cells (CD3D^+^, CD8A^-^, CCR7^low^, S100A4^int^, CD62L^high^, GZMA^-^) (79), T_TM_ cells (CD3D^+^, CD8A^-^, CCR7^low^, S100A4^high^, CD62L^low^, GZMA^+^, GZMB^-^, GZMH^-^, PRF1^-^, GNLY^+^) (80–82), T_EM_ cells (CD3D^+^, CD8A^-^, CCR7^-^, S100A4^high^, GZMA^+^, GZMB^+^, GZMH^+^, PRF1^+^ GNLY^+^) (83), T_EMRA_ cells (CD3D^+^, CD8A^-^, CCR7^-^, S100A4^high^, GZMA^+^, GZMB^high^, GZMH^+^, PRF1^high^, GNLY^high^, CCL4^+^) (40,84), T_REG_ cells (CD3D^+^, CD8A^-^, CCR7^low^, S100A4^high^, FOXP3^+^, IL2RA^+^, CTLA4^+^) (40,84), T_EX_ cells (CD3D^+^, CD8A^-^, CCR7^-^, GZMB^+^, PDCD1^+^, LAG3^+^) (85). All utilized code is deposited at https://github.com/GoffinetLab/HIV_scRNAseq-CD4-LRA-study. To visualize sequencing coverage of the viral genome in J1.1 T-cells, viral reads were extracted from the bam files of the CellRanger output. These were converted to bigwig files and visualised on tracks using bamCoverage (setting –normalizeUsing RPGC) and pyGenomeTracks from deeptools (86).

### Principal component analysis (PCA)

PCA was performed with the average expression values of all genes with detectable expression levels of mock-, Vorinostat- and Panobinostat-treated samples. Singular value decomposition (SVD) with imputations was used for calculation of the principal components using ClustVis (https://biit.cs.ut.ee/clustvis/) (87).

### Differentially expressed genes and pathway analysis

DEGs between individual treatments were identified using the 10x Genomics Loupe Browser (v. 5.0.1.); p-values were adjusted using the Benjamini-Hochberg correction for multiple testing. Pathway analysis was performed with the list of DEGs harbouring p-values <0.05, gene set enrichment analysis (GSEA) was performed using the Pathway Panther database (88–90). The results are described using the Normalised Enrichment Ratio (NER), the enrichment ratio indicates the degree of overrepresentation of a given gene set in the list of DEGs and is normalised to take different gene set sizes into account.

### Transcription factor activity analysis

Transcription factor activity analysis was performed using the dorothea R package following the guidelines for processing single cell RNA-seq data (91,92), with the following exception. The run_viper function from dorothea was altered to use the top 40,000 most variable genes, as determined by the FindVariableFeatures function from Seurat. Utilised code is available at https://github.com/GoffinetLab/HIV_scRNAseq-CD4-LRA-study.

### Module scores

The IFN signalling pathway (R-HSA-913531) and TCR signalling pathway (R-HSA-202403) gene sets from the Reactome database (93) were retrieved from the Molecular Signatures Database (MSigDB) (94). Cells were scored based on their expression of these genes using the AddModuleScore function in Seurat. They are referred to as the IFN signalling module and T-cell activation module scores as the pathways include genes canonically involved in response to IFN signalling and TCR-mediated T-cell activation, respectively.

### Flow cytometry

PBS-washed cells were immunostained for individual surface proteins using the following antibodies: anti-BST2/Tetherin-BV421 (#566381; BD), anti-CCR5/CD195-FITC (#555992; BD Biosciences), anti-CCR7/CD197-PE (#552176; BD Biosciences), anti-CD107a-BV421 (#562623; BD), anti-CD14-PE (#555398; BD), anti-CD16-FITC (#360716; Biolegend), anti-CD3-FITC (#561807; BD Biosciences), anti-CD4-APC (#555349; BD Biosciences), anti-CD25-APC (#340907; BD Biosciences), anti-CD45RA-FITC (#335039; BD Biosciences), anti-CD45RO-APC (#340438; BD Biosciences), anti-CD56-APC (#318310; Biolegend), anti-CD62L-FITC (#304804; Biolegend), anti-CD69-APC (#340560; BD Biosciences), anti-CXCR4/CD184-APC (#555976; BD Biosciences), anti-HLA-DR-FITC (#556643, BD Biosciences), anti-IL7R/CD127-PE (#557938; BD), anti-PD-1/CD279-PE (#21272794; ImmunoTools) and anti-TIM-3/CD366-FITC (#345022; Biolegend). For intracellular immunostaining, PBS-washed cells were PFA-fixed for 90 minutes and immunostained with the following antibodies diluted in 0.1% Triton X-100 in PBS: anti-HIV-1-core-antigen-FITC or -PE (#6604665 or #6604667; Beckman Coulter), rabbit-anti-human-MX1/2 (#sc-166412; Santa Cruz Biotechnology). A goat-anti-rabbit IgG conjugated to Alexa Fluor 647 (#A27040; Thermo Fisher) was used as a secondary antibody. Cell viability was analysed using the Dead Cell Apoptosis Kit for Flow Cytometry from Invivogen (#V13242), with early apoptotic cells scoring Annexin V-positive and late apoptotic cells scoring Annexin V- and propidium iodide (PI)-positive. Data acquisition and analysis was conducted using a FACS Celesta device (Becton Dickinson, Franklin Lakes, New Jersey, USA) with FlowJo (v.10.7.1).

### T-cell activation assays

CD4^+^ T-cells were isolated from healthy donors and pre-incubated with Vorinostat (500 nM), Panobinostat (50 nM), or left mock-treated (0.05% DMSO) for 16 hours followed by stimulation with a peptide pool of a broad range of HLA-subtypes and infectious agents (CEFX Ultra SuperStim Pool; JPT) (1 µg/ml) and 1 μg/ml mouse-anti-human CD28 antibody (#556620; BD Biosciences), or IL-2 (20 IU/ml) and PHA (1 µg/ml). T-cell activation and exhaustion was quantified by flow cytometry 48 hours post-stimulation.

### HIV-1 reactivation assays in J1.1 T-cells

J1.1 T-cells were PBS-washed, resuspended in supplemented RPMI and incubated with Panobinostat (50 nM), Vorinostat, (5000 nM), Romidepsin (50 nM), Bryostatin (20 nM), JQ1 (5000 nM) or the corresponding volume of DMSO, if not otherwise stated for 48 hours. In some assays, T-cells were co-treated or pre-treated for 24 hours with individual IFNs at indicated concentrations.

### Quantitative RT-Q-PCR

Total RNA extraction using the Direct-zol RNA extraction kit (Zymo) including DNase treatment to remove residual DNA contaminations, was followed by cDNA synthesis (NEB, Invitrogen) and quantification of relative mRNA levels using a LightCycler 480 Instrument II (Roche) and Taq-Man PCR technology. For human *IFIT1* and *MX2*, a premade primer-probe kit was purchased from Applied Biosystems (Assay ID: Hs01911452_s1; Hs01550813_m1; respectively. Relative mRNA levels were determined in multiplex reactions using the ΔΔCt method and human *RNASEP* (#4316844; Applied Biosystems) as internal reference. Data analysis was performed using LightCycler Software 4.1 (Roche).

### Immunoblotting

For immunoblotting, cells were lysed using the M-PER mammalian protein extraction reagent (Thermo Fisher Scientific) following manufacturer’s recommendations. Cell cultures supernatants were centrifuged over a 20% sucrose cushion at 20,000 x g, for 60 mins at 4°C, resuspended in 1x SDS buffer at heat-inactivated for 10 mins at 95°C. Samples were run on a 10% SDS-PAGE, transferred onto nitrocellulose membranes using a semi-dry transfer system (Bio-Rad Laboratories) and membranes were blocked with 5% milk in TBS for one hour before incubation with the primary antibody at 4°C overnight. The following primary antibodies were used: anti-90K/LGALS3BP (#AF2226: R&D Systems), anti-BST2/Tetherin (#390719; Santa Cruz), anti-ERK2/MAPK (#sc-153; Santa Cruz) anti-GAPDH (#NB300-221, Novus Biologicals), anti-HIV-1 p24 (#11-327; ExBio), anti-HIV-1 gp120 (provided by Valerie Bosch), anti-IFIT1 (#TA500948; OriGene), anti-IFITM-3 (#AP1153a; Abgent), anti-ISG15 (#166755; Santa Cruz) and anti-MX2 (#sc166412; Santa Cruz). Secondary antibodies conjugated to Alexa680 or Alexa800 fluorescent dyes were used for detection and quantification by the Odyssey Infrared Imaging System (LI-COR Biosciences).

Release of HIV-1 p24 capsid was quantified by detecting p24 as described above in supernatant and cell samples from corresponding cell cultures and dividing the p24 signal intensity from supernatants by the sum of p24 intensities from both fractions.

### Tzm-bl HIV-1 infectivity assay

To quantify infectivity released in supernatant, 30,000 Tzm-bl cells per 96-well were pre-treated with 10 μM of the JAK/STAT inhibitor Ruxolitinib for 16 hours to minimise the influence of potentially transferred IFNs from the tested supernatants. Then, cells were infected with HIV-1 for 48 hours before luminometric quantification of luciferase activity using the Luciferase Assay System (Promega).

### HIV-1 Env cell surface staining

To quantify HIV-1 Env cell surface expression, broadly neutralising antibodies 3BNC117 (ARP-12474), 10-1074 (ARP-12477) and PG16 (ARP-12150) were obtained from the NIH HIV Reagents Program. PBS-washed cells were immunostained with a cocktail of 5 μg/ml of each antibody for 30 mins at 4°C, followed by PBS washing and immunostaining with secondary AlexaFluor633 anti-human antibodies (Thermo Fisher Scientific, #A-21091) for 30 mins at 4°C in the dark. Samples were PBS-washed and fixed with 4% PFA for 90 minutes. Samples were acquired and analysed on a FACS Celesta machine (BD Biosciences) and FlowJo Software (V.10.7.1), respectively.

### Thin section electron microscopy

Infected T-cells were fixed with 2.5% glutaraldehyde in 0.05 M Hepes buffer (pH:7.2) and incubated at room temperature for two hours. Afterwards, µ-Dishes were filled with the fixative buffer and cells were embedded in the chambers by using Epon resin (protocol with tannic acid and uranyl acetate block contrasting (95). Thin sections (60-70 nm thick) were produced with an ultramicrotome, contrasted with uranyl acetate and lead citrate and investigated with a transmission electron microscope (JEM-2100, Jeol) operated at 200 kV. Images were recorded using a side-mounted CCD camera (Veleta, EMSIS) with 2048×2048 pixels.

### Antibody-dependent cellular cytotoxicity (ADCC) assay

ADCC assays were adapted from previous reports (96). Briefly, J1.1 T-cells were treated as described above for 48 hours, before washing and staining with green CellTracker (#C2925, Thermo Fisher Scientific), following manufacturer’s recommendations. Labelled J1.1 T-cells were then co-cultured with freshly isolated PBMCs (or NK cells or monocytes where indicated) at a 1:1 ratio in a 96-well U-bottom plate in the presence of indicated dilutions of serum from PLHIV (ARP-3957; obtained from the NIH HIV Reagents Program). A separate well without serum was set up for each co-culture condition in parallel. The plates were then centrifuged for one min, 300 x *g* and then incubated at 37°C, 5% CO_2_. After four hours, co-cultures were PBS-washed, fixed with 4% PFA, immunostained for intracellular HIV-1 p24 expression and p24 CA-positive J1.1 T-cells were quantified by flow cytometry. Percent of ADCC was calculated using this formula (see also **Sup. Fig. 8**):

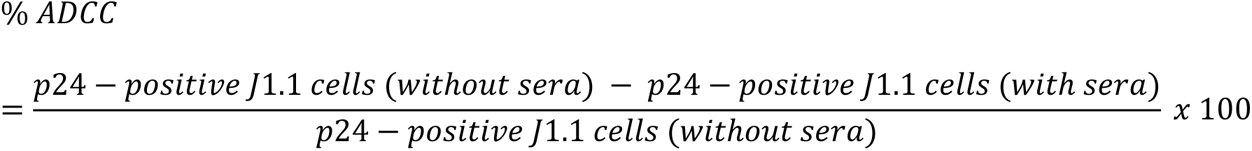

### NK cell activation and CD107a degranulation assay

NK cell activation and degranulation activity in PBMCs co-cultured with reactivated J1.1 T-cells was analysed through CD16 and CD107a immunostaining of NK cells. ADCC assays were performed as described above with the following modifications (97): Co-cultures were supplemented with a 1:20 dilution of anti-CD107a-BV421 antibodies. After one hour of incubation, a final concentration of 10 μg/ml Brefeldin A and 6 μg/ml Golgi-Stop (BD Biosciences) were added, preventing the exocytosis of cytokine-containing vesicles, and the acidification of endocytic vesicles and CD107a protein degradation, respectively. Co-cultures were incubated for another three hours at 37°C, 5% CO_2_, before immunostaining for CD16 and CD56 cell surface expression. CD16 and CD107a expression was assessed by flow cytometry specifically in CD56-positive NK cell populations (see also **Sup Fig. 9**).

### Live-cell imaging of J1.1 T-cell/NK cell co-cultures

To analyse the dynamics of NK cell-mediated ADCC against reactivating J1.1 T-cells, live cell imaging of indicated co-cultures was performed. J1.1 T-cell cultures treated with Panobinostat or Panobinostat/IFN-α2a for 48 hours were individually labelled with CellTracker^TM^ Green-CFMDA or CellTracker^TM^ Deep Red CellTracker (both Thermo Fisher Scientifics). Panobinostat-treated, Panobinostat/IFN-α2a-treated J1.1 T-cells and NK cells were co-cultured at a 1:1:1 ratio in the presence of 1:100 dilution of sera from PLHIV and the LIVE/DEAD Aqua Dead Cell dye (Thermo Fisher Scientific, #L34965) to assess dying cells. The dead cell dye reacts with free amids in the cellular cytosol, thus specifically staining cells with porous membranes as an early surrogate of cell death. Co-cultures were monitored with a Zeiss LSM800 Airyscan Confocal Microscope in a 37°C, 5% CO_2_ atmosphere with images being acquired every three minutes for a total of four hours. Images were analysed and merged using Zeiss ZEN Blue (V3.3).

### Data presentation and statistical analysis

Graphs and figures were generated using Graphpad Prism 9 (v.9.5.1) and Adobe Illustrator 2021. Heatmaps were generated using the ClustVis web tool (https://biit.cs.ut.ee/clustvis/) with unit variance scaling for rows (87). If not otherwise stated, bars or circles represent the mean of the indicated number of experiments and error bars indicate the S.E.M. Statistical significance for paired data sets were tested using Graphpad Prism 9 and paired student’s t-testing, p-values ≤ 0.05 were considered significant and displayed in the figures as follows: p < 0.05 *; p < 0.01 ** or p < 0.001 ***. p-values ≥ 0.05 were considered not significant (n.s.). Pearson’s correlation analyses were performed for the indicated datasets with pearson correlation coefficients and two-tailed p-values being displayed.

### Data availability

The single cell RNA-sequencing data will be deposited at the Gene Expression Omnibus (GEO) database (accession number #).

## Supporting information

Prigann et al., Supplemental Material

Prigrann et al., Supplemental Video 1

## ACKNOWLEDGEMENTS

We thank the PLHIV who agreed to participate in the Seroconverter Study and the uninfected donors to provide their blood for research. We thank Valerie Bosch and the NIH AIDS Research & Reference Reagent Program for providing essential reagents. We thank the Genomics Platform of the Berlin Institute of Health for next-generation sequencing. This work was supported by funding to C.G. by Berlin Institute of Health (BIH); by Hector Foundation, project M2101; by DFG priority program 1923 “Innate Sensing and Restriction of Retroviruses,” grant GO2153/4); by DZIF, TTU HIV, grant 04.820 to C.G. and by a Professorship Award (APR9/1017) of the Academy of Medical Sciences awarded to C.G. We thank Christian Drosten for constant support. A.P. is supported by the Brazilian Federal Foundation for Support and Evaluation of Graduate Education, CAPES. J.K. is supported by the Center of Infection Biology and Immunity (ZIBI) and Charité PhD Program.

## AUTHOR CONTRIBUTIONS

JP, AJP, JJ, LM, JF, TS, performed experiments.

JP, DP, AJP, EW, LM, JF, TS, CF, SV, MS analysed data.

UK, BGB, KM, NB, LL, AT, KS provided essential resources.

AT, SS, UD, MiL, NB, MaL, CG supervised.

CG acquired funding and managed the project.

JP, DP, CG wrote the manuscript.

