## Supplementary material for "Type I interferons sensitise HIV-1-reactivating T-cells for NK cell-mediated elimination despite HDACi-imposed dysregulation of innate immunity": Prigann et al., Supplemental Material

### SUPPLEMENTARY MATERIALS

**Supplementary Table 1.**

**Supplementary Figure 1-11**

**Supplementary Movie 1.**

**Prigann et al., Supplementary Table 1**

| Donor ID | Seroconversion | Treated since | Viral Load (viral copies/ml) | CD4 counts (cells/µl) |
|---|---|---|---|---|
| 1 | 2006 | 2008 | < 50 | 1048 |
| 2 | 2008 | 2011 | < 50 | 526 |
| 3 | 2017 | 2018 | < 50 | 629 |

**Supplementary Table 1. Clinical data of the three PLHIV.**

Shown are the dates of infection and ART initiation as well as viral loads and counts of $CD4^+$ T-cells in the peripheral blood. The detection limit for viral copies/ml is 50; samples with undetectable levels are indicated with “< 50”.

Prigann ***et al.***, **Supplementary Figure 1**

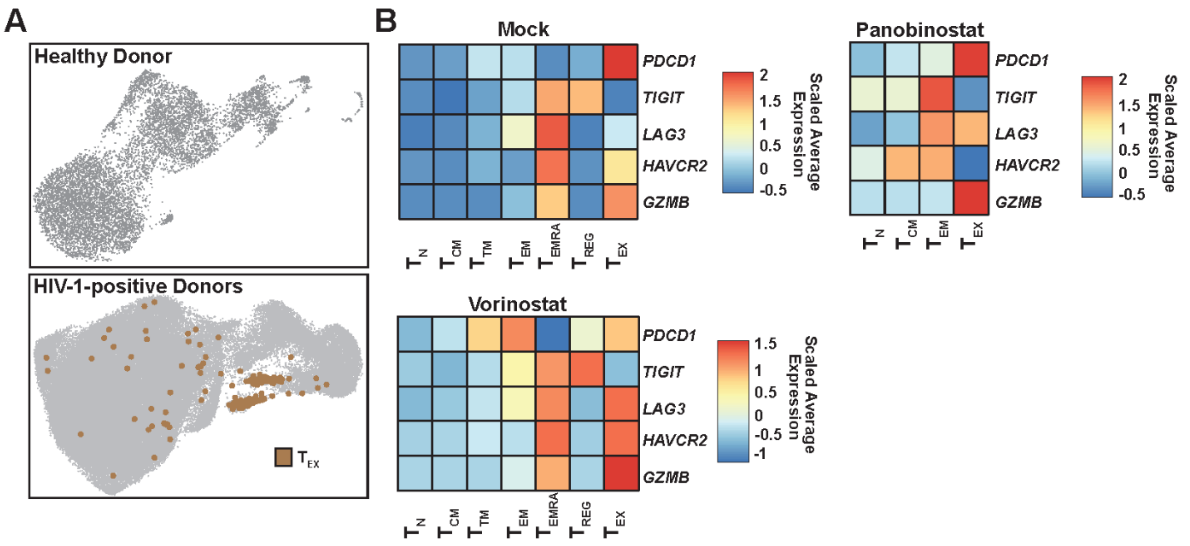


**Supplementary Figure 1. A subset of CD4⁺ T-cells, identified specifically in cells isolated from PLHIV, resembles $T_{EMRA}$ cells, with additional expression of the exhaustion marker *PDCD1***

(**A**) UMAP plots of CD4⁺ T-cells from an HIV-1-negative donor (n=1) and from aviremic PLHIV (n=3). Exhausted T-cells are coloured in brown.

(**B**) Heatmaps showing the scaled average expression of T-cell exhaustion markers, *PDCD1*, *TIGIT1*, *LAG3*, *HAVCR2*, *GZMB* in the different CD4⁺ T-cell subsets of PLHIV (n=3) upon mock-treatment or treatment with Vorinostat or Panobinostat.

**Prigann *et al.*, Supplementary Figure 2**

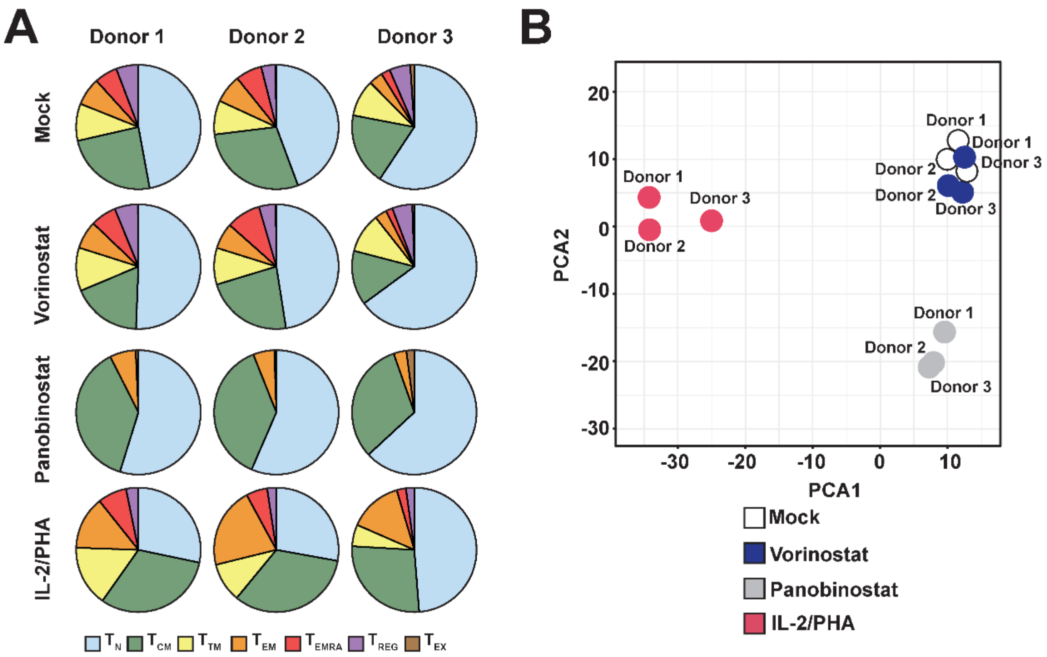


**Supplementary Figure 2. CD4⁺ T-cell subset distribution and the transcriptomic landscape is determined by treatment condition in a donor-independent manner**

(**A**) Pie charts displaying the CD4⁺ T-cell subset distribution of samples obtained from the individual PLHIV after indicated treatment.

(**B**) PCA based on the global gene expression profile of the individual samples after indicated treatment.

**Prigann *et al.*, Supplementary Figure 3**

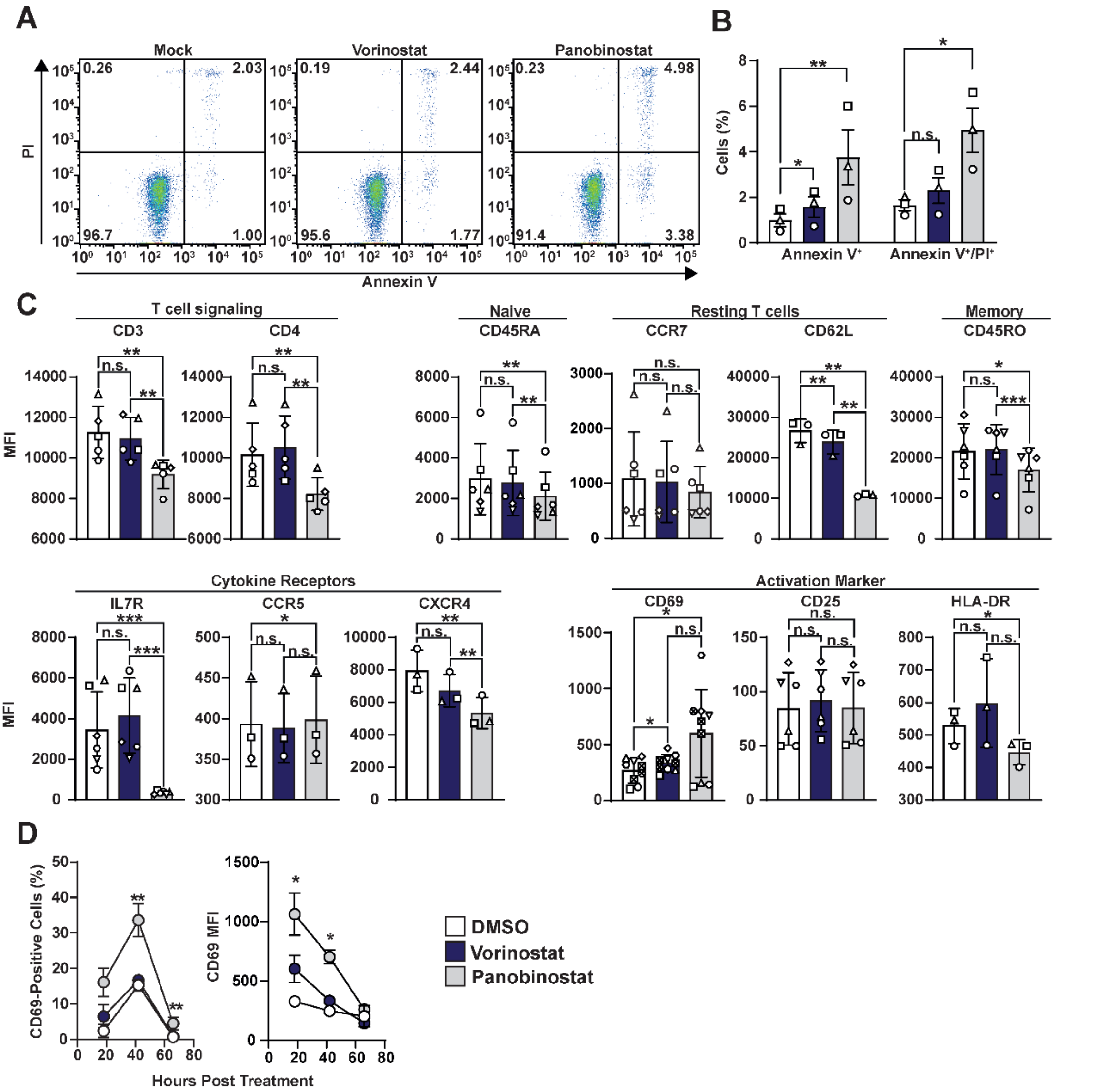


**Supplementary Figure 3. Panobinostat treatment modulates expression of CD4⁺ T-cell markers**

CD4⁺ T-cells from HIV-1-negative donors were treated with 0.5% DMSO (mock), Vorinostat (500 nM ) or Panobinostat (50 nM).

(**A**) Representative Annexin V/PI cell apoptosis staining upon indicated treatment for 48 hours.

(**B**) Percentage of Annexin V⁺- and Annexin V⁺/PI⁺-positive CD4⁺ T-cells from three individual donors upon indicated treatment for 48 hours.

(**C**) Mean fluorescence intensity (MFI) of cell surface expression of indicated proteins upon indicated treatment for 48 hours.

(**D**) Percentage of $CD69^+$ T-cells and MFI of CD69 expression upon indicated treatment of cells from three donors.

**Prigann *et al.*, Supplementary Figure 4**

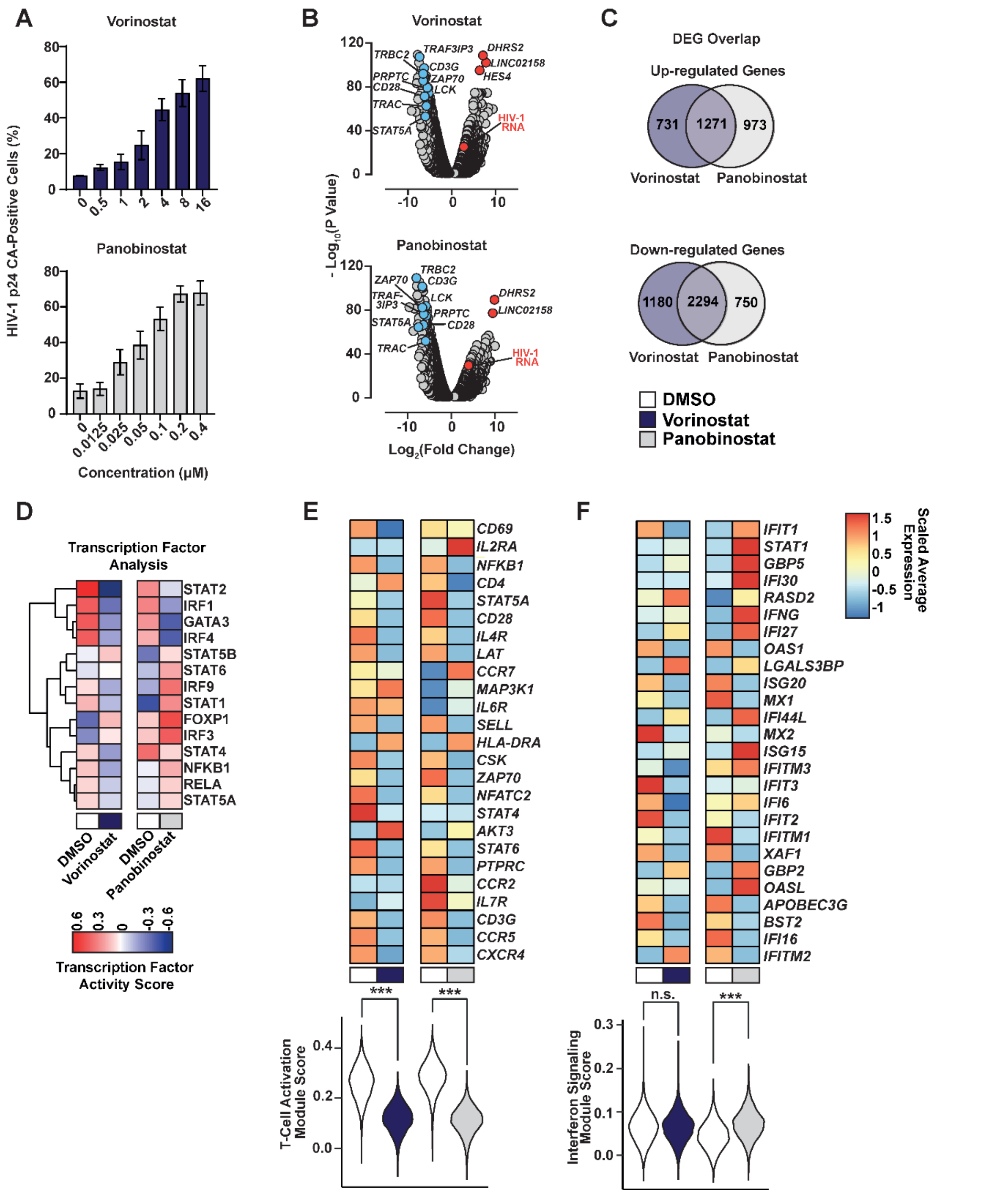


**Supplementary Figure 4. HDACi-induced transcriptome in the J1.1 T-cell model of HIV-1 latency**

(**A**) J1.1 T-cells were treated with the indicated concentration of Vorinostat or Panobinostat for 40 hours before quantification of intracellular HIV-1 p24 CA expression by flow cytometry.

(**B**) - (**F**) J1.1 T-cells were treated with 16 µM Vorinostat, 1.6% DMSO (DMSO I), 200 nM Panobinostat or 0.2% DMSO (DMSO II) for 40 hours prior to single cell RNA-sequencing analysis.

(**B**) Volcano plot showing significant DEGs of Vorinostat- or Panobinostat-treated samples compared to mock samples.

(**C**) Overlap of genes up- and down-regulated after Vorinostat- or Panobinostat-treatment compared to mock-treatment.

(**D**) Activity analysis of selected transcription factors.

(**E**) Heatmap showing expression of selected genes involved in T-cell activation and signalling and T-cell activation module scores. Statistical significance for the module score was tested using Wilcoxon signed-rank testing.

(**F**) Heatmap showing expression of selected ISGs and IFN signalling module scores. Statistical significance for the module score was tested using Wilcoxon signed-rank testing.

**Prigann *et al.*, Supplementary Figure 5**

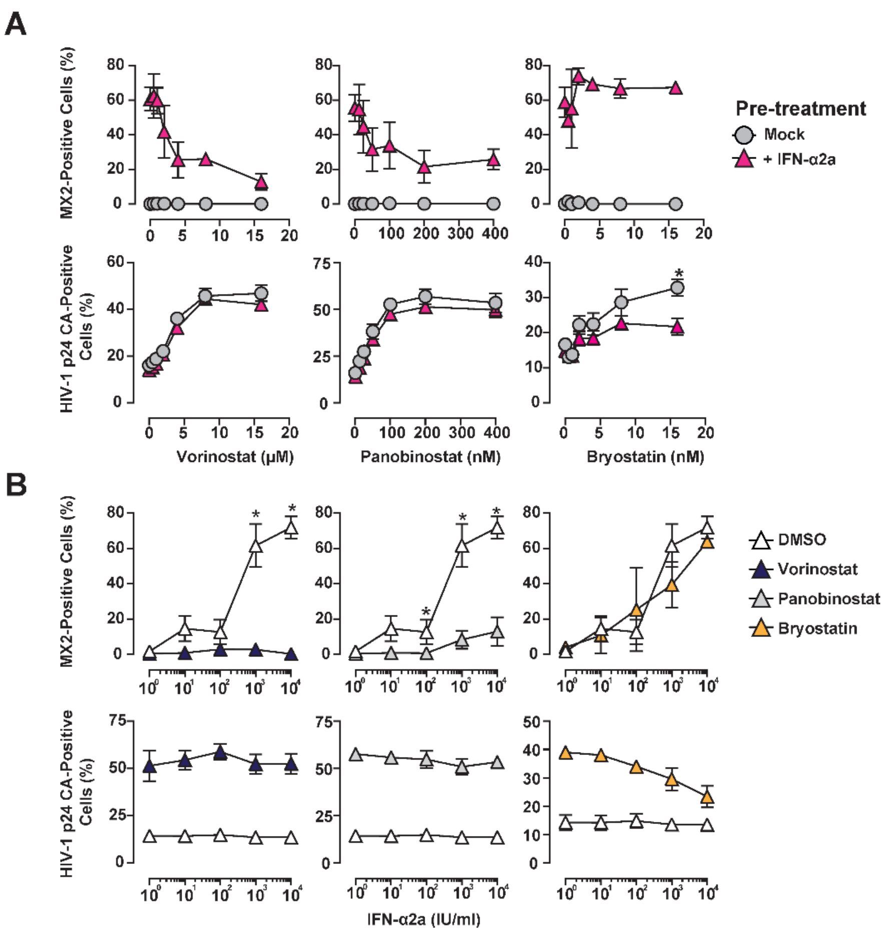


**Supplementary Figure 5. HDAC inhibition suppresses type I IFN-induced ISG expression**

(**A**) J1.1 T-cells were pre-treated with IFN-α2a (500 IU/ml) or mock-treated for 24 hours before incubation with indicated concentrations of Panobinostat, Vorinostat, Bryostatin or DMSO. Intracellular HIV-1 p24-CA and MX2 expression was quantified 40 hours post addition of LRAs by flow cytometry.

(**B**) J1.1 T-cells were cultured in the presence of Panobinostat (200 nM), Vorinostat (8 µM), Bryostatin (10 nM) or DMSO in combination with indicated concentrations of IFN-α2a. Intracellular HIV-1 p24 CA and MX2 expression was quantified after 40 hours.

Experiments were performed as 3-4 independent replicates.

**Prigann et al., Supplementary Figure 6**

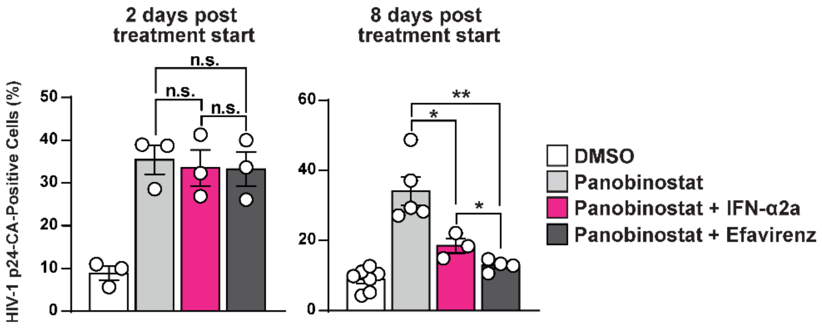


**Supplementary Figure 6. HIV-1 reverse transcriptase inhibitor Efavirenz blocks spread of HIV-1 upon Panobinostat-triggered HIV-1 reactivation.**

J1.1 cells were treated with 50 nM Panobinostat alone, in combination with either 100 IU/ml IFN-α2a or 100 nM Efavirenz or mock-treated with DMSO for 2 or 8 days before analysing HIV-1 p24 CA-positive cells by flow cytometry.

**Prigann et al., Supplementary Figure 7**

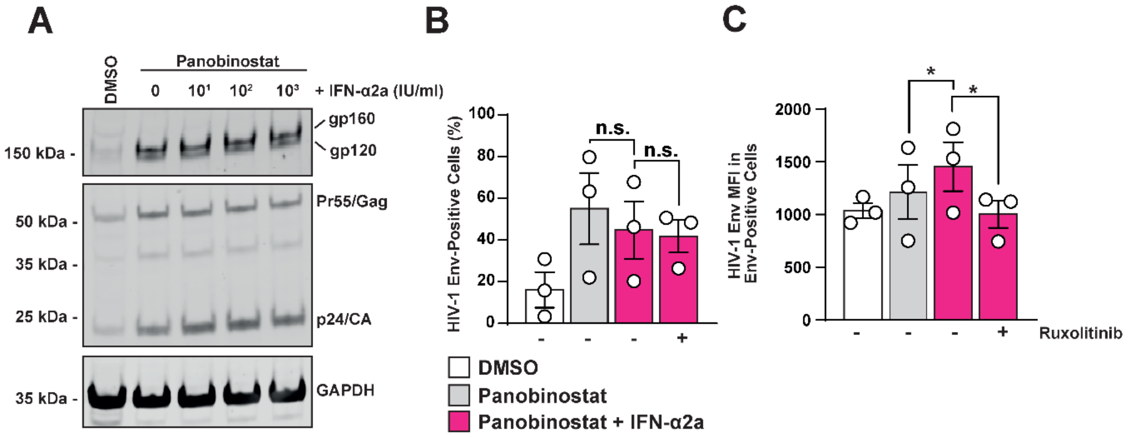


**Supplementary Figure 7. Influence of IFN-α2a signalling on HIV-1 gp160 processing and Env cell surface expression.**

(**A**) J1.1 T-cells were treated with DMSO, 50 nM Panobinostat alone or in combination with indicated concentration of IFN-α2a for 48 hours and analysed for HIV-1 gp160, gp120, Pr55Gag and p24 CA protein expression and processing via immunoblotting.

(**B-C**) J1.1 T-cells were treated 50 nM Panobinostat in the absence or presence of 100 IU/ml IFN-α2a and 10 μM Ruxolitinib, immunostained for surface HIV-1 Env and analysed for (**B**) percentage of HIV-1 Env-positive cells or (**C**) Env MFI in Env-positive cells.

**Prigann et al., Supplementary Figure 8**

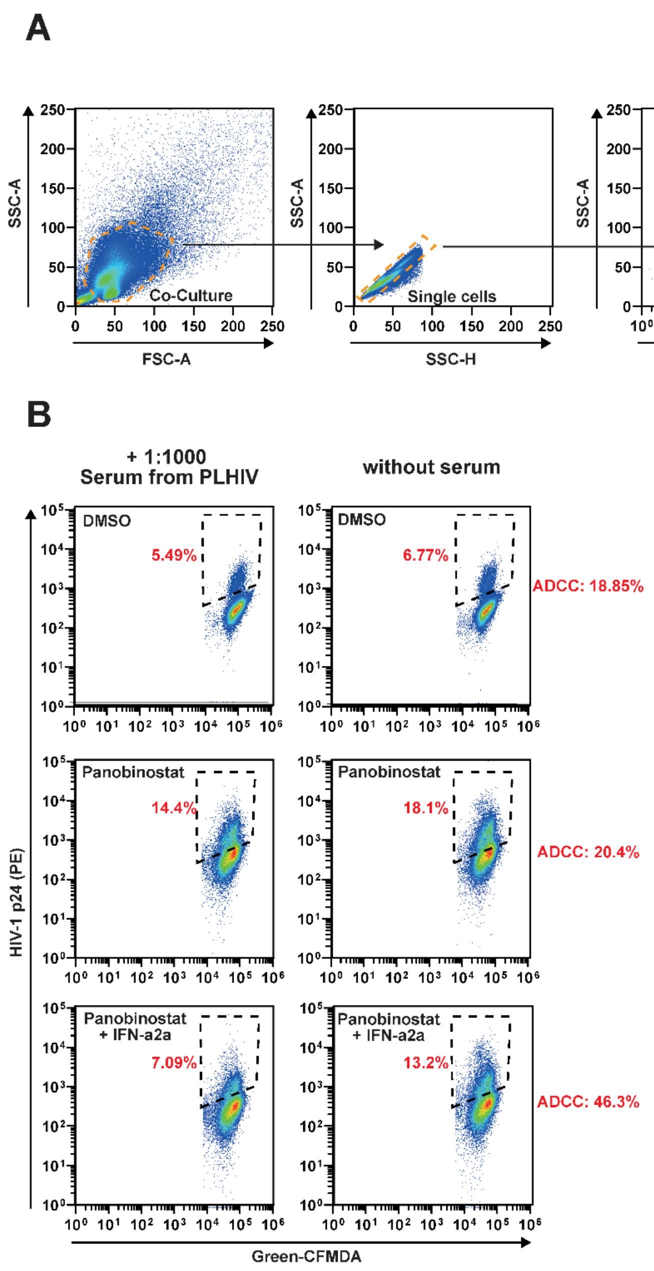


**Supplementary Figure 8. Gating strategy and analysis of the ADCC assay**

To characterise ADCC targeting reactivated J1.1 T-cells, freshly isolated PBMCs were co-cultured with cell-tracker Green-CFMDA stained J1.1 T-cells at a 1:1 ratio for four hours. The co-culture was then subjected to intracellular HIV-1 p24 CA-PE immunostaining.

(**A**) The gating strategy to determine the percentage of p24 CA-positive J1.1 T- cells in the co-culture is shown.

(**B**) A representative dataset is shown for which the ADCC percentage was calculated.

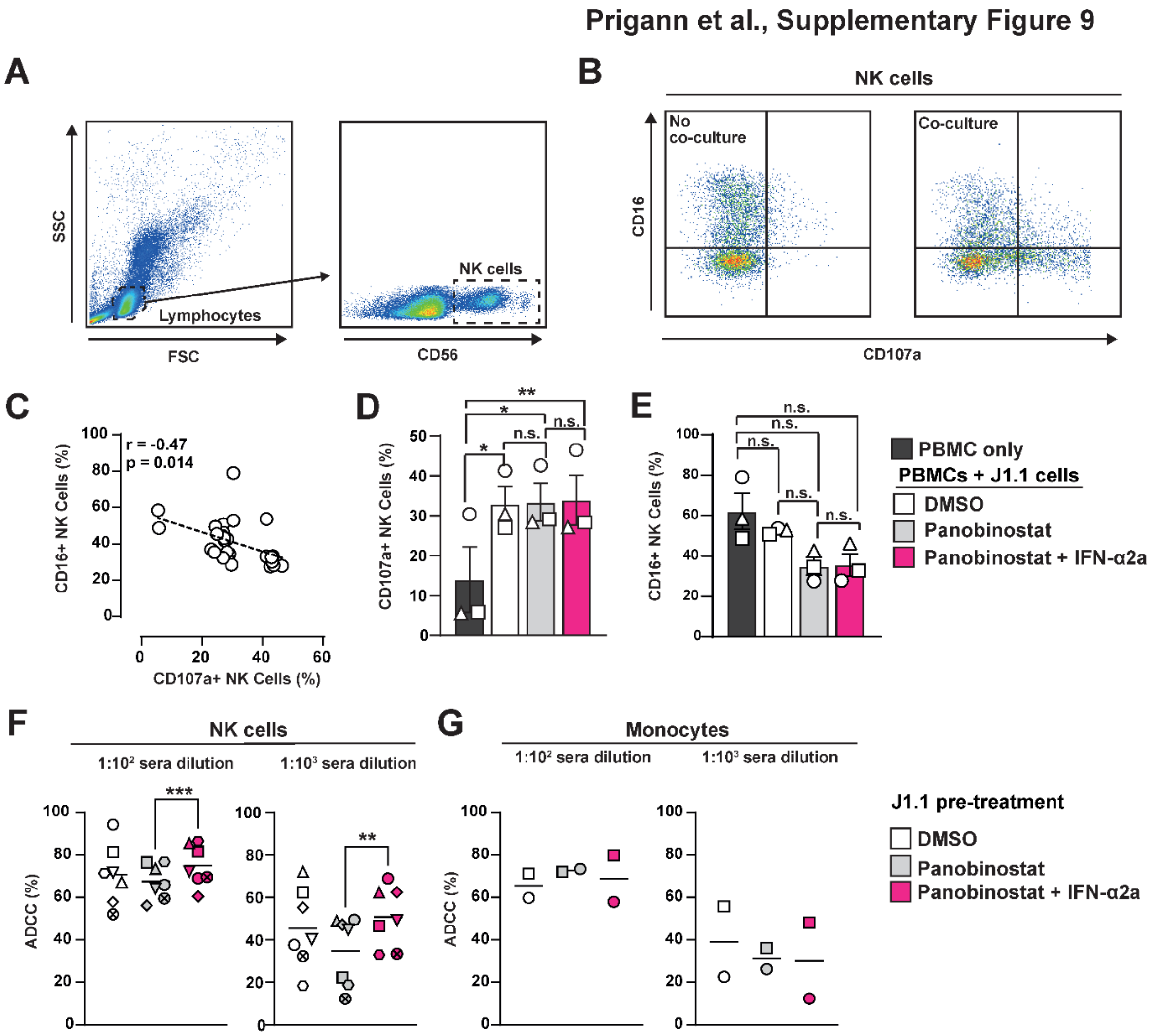


**Supplementary Figure 9. NK cells display signs of degranulation and cellular activation upon co-culturing with J1.1 T-cells in an IFN-independent manner.**

PBMC co-cultures with J1.1 T-cells were established as described before, but complemented with anti-CD107a-BV421 antibodies for the duration of the co-culture. After one hour of co-culture, Brefeldin A and Golgi Stop were added and after a total of four hours of co-culture, cells were immunostained for CD56 and CD16 cell surface expression.

(**A**) Representative plots showing the gating strategy to identify NK cells in the co-culture.

(**B**) Representative plots showing CD16- and CD107a-positive NK cells within PBMCs alone or in co-culture with J1.1 T-cells.

(**C**) Pearson's correlation analysis of the percentage of CD16- and CD107a-positive NK cells combining the data of all analysed co-cultures.

(**D**) Quantification of CD107a-positive cells within the NK cell populations following culturing PBMCs as indicated.

(**E**) Quantification of CD16-positive cells within the NK cell populations following culturing PBMCs as indicated.

(**F**) - (**G**) J1.1 T-cells were treated with DMSO, 50 nM Panobinostat alone or in combination with 100 IU/ml IFN-α2a for 48 hours before co-culturing with isolated NK cells (**F**) or monocytes (**G**) for four hours in the presence of serum from PLHIV. The percentage of ADCC was calculated as described before. Symbols indicate individual experimental replicates.

**Prigann et al., Supplementary Figure 10**

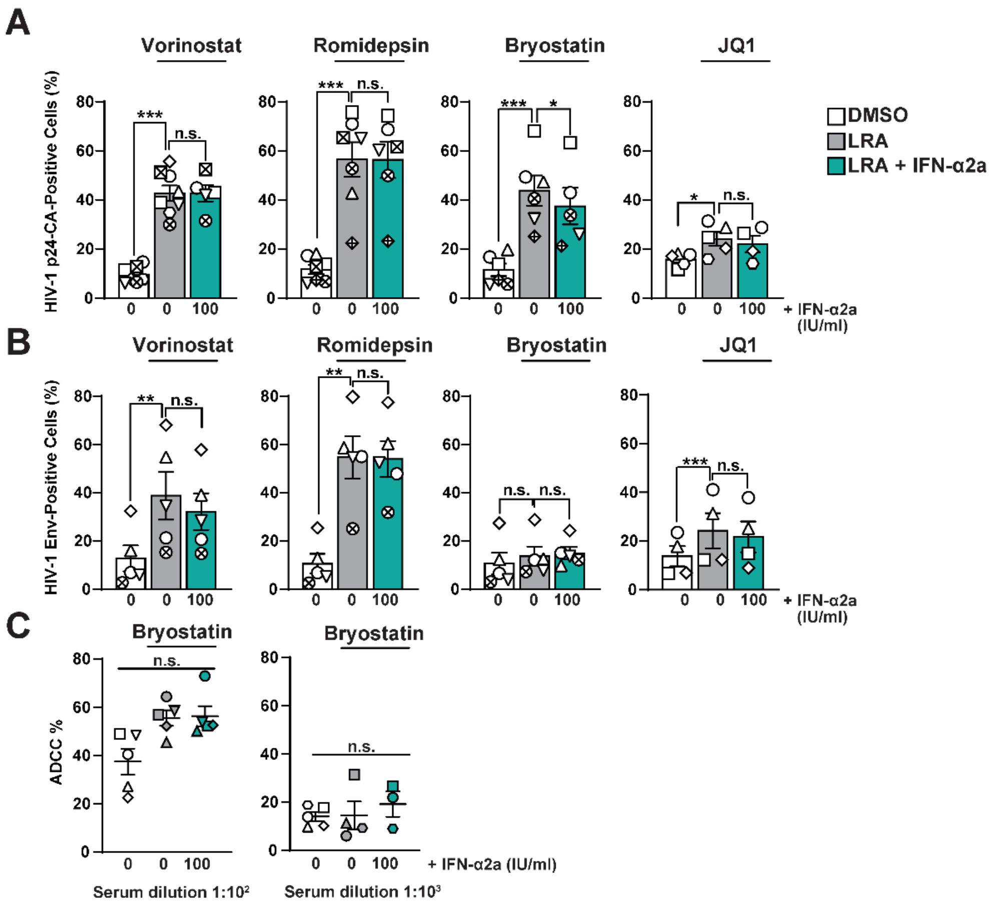


**Supplementary Figure 10. IFN-α2a-enhanced HIV-1 Env cell surface accumulation and ADCC sensitization occur in the context of HDACi-, but not PKC agonist-mediated HIV-1 reactivation**

J1.1 T-cells were treated with 4 µM Vorinostat, 25 nM Romidepsin, 20 nM Bryostatin or 10 µM JQ1 in the presence or absence of 100 IU/ml IFN-α2a or DMSO for 48 hours.

(**A**) Quantification of the percentage HIV-1 p24 CA-positive cells by flow cytometry.

(**B**) Quantification of HIV-1 Env cell surface-positive cells by flow cytometry.

(**C**) Bryostatin-treated cells were subjected to ADCC assays, as described before, using 1:100 and 1:1000 dilutions of serum from PLHIV

**Prigann et al., Supplementary Figure 11**

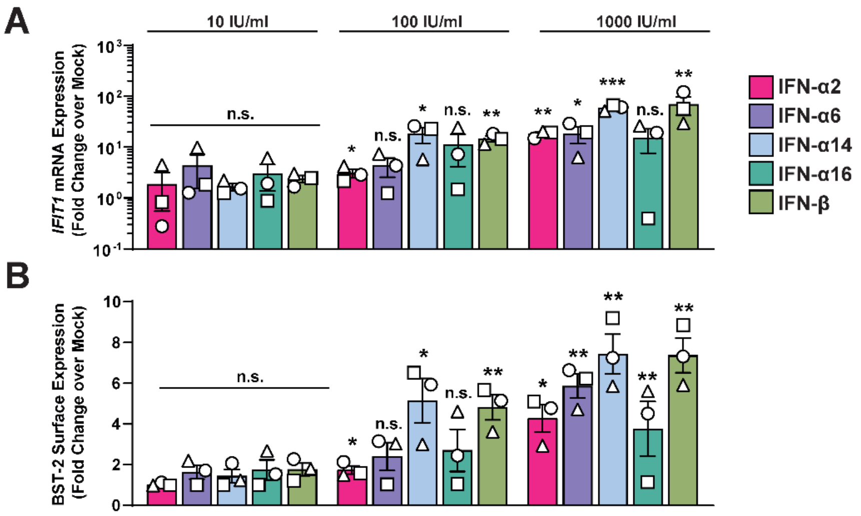


**Supplementary Figure 11. IFN-α14 and IFN-β exhibit most pronounced ability to induce ISG expression in latently HIV-1-infected J1.1 T-cells**

J1.1 T-cells were treated with indicated concentrations of type I IFNs for 72 hours and analysed for (**A**) *IFIT1* mRNA expression via RT-Q-PCR and (**B**) BST-2/Tetherin cell surface expression via flow cytometry. Bars show the mean of three independent experiments.

**Supplementary Movie 1. Addition of IFN-α2a to Panobinostat results in accelerated killing of HIV-1-reactivating T-cells.**

J1.1 T-cells were treated with 50 nM Panobinostat alone or in combination with 100 IU/ml IFN-α2a for 48 hours and individually stained with CellTracker™ Green-CFMDA and Deep Red, respectively. Stained J1.1 T-cells were co-cultured with non-stained, freshly isolated NK cells at a 1:1:1 ratio in the presence of a cell death marker (blue) and a 1:100 dilution of sera from PLHIV containing anti-HIV-1 Env antibodies. The co-culture was monitored using live-cell imaging for four hours with images acquired every three minutes.
